# Subthalamic Signature of Freezing of Gait in Parkinson’s Disease

**DOI:** 10.1101/2025.01.10.632393

**Authors:** Antoine Collomb-Clerc, Mathieu Yeche, Adèle Demain, Angèle Van Hamme, Claire Olivier, Hayat Belaid, Déborah Ziri, Stéphane Derrey, Sara Fernandez-Vidal, Katia Lehongre, Carine Karachi, Brian Lau, Marie-Laure Welter

## Abstract

Freezing of gait (FOG) is a significant disability in Parkinson’s disease (PD). Deep brain stimulation (DBS) of the subthalamic nucleus (STN) only partially alleviates it, with approximately one-third of patients experiencing worsening FOG within a year after surgery. The precise role of STN dysfunction in gait disabilities and FOG remains not fully elucidated. To investigate this, we recorded gait and STN local field potential (LFP) activity in 38 PD patients, both Off and On dopamine medication. Our analysis focused on the relationship between gait performance and STN neuronal activity, particularly examining differences in LFP activity across the posterior-sensorimotor and central-associative regions of the STN. When Off dopamine medication, 12 patients experienced FOG during recordings, with a total of 263 FOG episodes documented. Even in trials without FOG episodes, these patients exhibited altered gait initiation strategies, prioritizing stepping rhythm to manage balance and initiate walking. In contrast, non-FOG patients maintained a higher walking pace. STN activity patterns revealed key differences. In FOG patients, weaker STN alpha/low beta band activity in the STN was associated with walking pace, while stronger decreased low beta band activity correlated with rhythm and balance control. This low beta band association extended from the posterior-sensorimotor to the central-associative STN. In contrast, non-FOG patients showed a more restricted relationship between low beta band activity and gait performance, confined to the posterior STN. As stepping rhythm deteriorated further in FOG patients, FOG episodes occurred. FOG episodes were preceeded by a significant positive relationship between high beta power and rhythm restricted to the posterior STN, with a reverse negative relationship with pace, and a disruption in low beta desynchronization across both posterior and central STN regions. Dopamine medication significantly improved gait patterns, and partially restored STN neuronal activity, reducing differences between FOG and non-FOG patients. These findings differentiate two FOG states, i.e. predisposition and occurrence, each associated with distinct gait initiation strategies and STN activity patterns. They suggest distinct pathophysiological roles of low and high beta band STN activity within specific STN regions in regulating gait and FOG. These findings provide key insights for refining targeted DBS therapies.

## Introduction

Freezing of gait (FOG) – a sudden inability to lift the feet during walking – is a severe motor disability in Parkinson’s disease (PD)^1^. FOG is episodic and typically occurs during gait initiation, turning, navigating tight spaces, or during multitasking scenarios like carrying objects or talking while walking^1^. Affecting 50–70% of PD patients after a decade of disease progression, FOG severely impacts mobility, increases the risk of falls, morbidity, mortality, and healthcare costs. FOG is inconsistently improved by dopaminergic therapy, worsens over time, and leaves patients with limited treatment options^2^.

Deep brain stimulation (DBS) of the subthalamic nucleus (STN) effectively improves parkinsonian motor signs, but its effects on gait and FOG vary. Approximately one-third of patients experience worsened or new-onset FOG within the year following surgery^3,4^. This variability may be linked to physiological and anatomical differences between patients. Beta-band (13-35 Hz) activity measured within the STN, and the relationship between this activity and motor/premotor cortical activity has also been extensively studied in the STN of PD patients at rest^5–7^. Exaggerated beta activity with prolonged bursts is likely pathological as its reduction by dopaminergic treatment or STN-DBS^8–10^ correlates with improvement of parkinsonian motor signs^11,12^. During stepping in place, alternation between low (13-20 Hz) and high (21-35 Hz) beta band activity has been observed^13,14^. PD patients with FOG exhibit increased STN beta activity at rest^15^, while stepping in place^10^, and during virtual^16,17^ or real walking tasks^18–25^. Elevated theta and low beta band activity have also been linked to FOG vulnerability during single– and dual-task walking^25,26^ and bicycling^24^. A loss of movement-related alpha and beta desynchronization, along with low-frequency cortico-STN decoupling, has also been reported preceding and during FOG episodes.^25,27^ Anatomically, beta oscillaroty activity was reported to be predominantly located within the posterior-dorsal STN,^28^ and associated with bradykinesia and rigidity.^12^ However, some studies suggest that high-beta band activity is more widely distributed within the STN,^28,29^ and a stronger cross-frequency coupling between high-frequency oscillations and beta band activity has been observed in the more inferior-anterior region of the STN in PD patients with postural instability and gait difficulty (PIGD)^29^. With directional STN-DBS, targeting the anterior-central STN region – as opposed to the posterior-dorsal STN – yields better outcomes for gait disorders and FOG^30^, with recruitment of fiber pathways connected to the premotor and sensorimotor cortices^31,32^, as well as the mesencephalic locomotor region (MLR)^31^. Together, these findings underscore the critical role of STN dysfuntion in gait and postural disabilities in PD patients, including FOG ^10,21^. They also highlight that these abnormalities are likely unevenly distributed within the STN, depending on the severity of gait and balance disorders. These insights emphasize the need to deepen our understanding of STN neural dynamics to enhance FOG treatment and guide the development of adaptive DBS strategies^33,34^.

Here, we investigated the specific link between STN neuronal activity recorded in the posterior and central STN areas, and gait/postural abilities, including FOG, in 38 PD patients. We performed simultaneous recordings of STN local field potentials (LFPs) and gait, with precise determination of FOG events. We hypothesized that beta band activity would differ based on FOG predisposition and/or occurrence^21,26^, and that pathological beta modulation would exhibit differential distributed within the STN^3,28^.

## Materials and methods

### Participants

We recruited thirty-eight people with Parkinson’s disease (**Supplementary Table 1**) scheduled to receive deep brain stimulation of the subthalamic nucleus (STN-DBS) at the Pitié-Salpêtrière Hospital and Rouen University, in two clinical research trials, between February 2013 and September 2021 (GB-MOV study, INSERM promotion and MAGIC study, CHU ROUEN promotion)^3^. For both trials, the inclusion criteria were: 1) diagnosis of PD based on the UK brain bank criteria; 2) age between 18 and 70 yrs; 3) validation of the indication of STN DBS according to the local neurosurgical and neurological staff{Citation}, including high responsiveness of motor disability to levodopa treatment, existence of levodopa related motor complications, no ongoing psychiatric disorders or dementia (Mini-Mental Status score greater than 24), and no contraindication to surgical intervention for DBS implantation; 4) voluntarily and informedly agreed to participate in the study and signed a written consent and 5) had social insurance. For the MAGIC study, additional inclusion criteria were: 1) freezing of gait in the DOPA^OFF^ condition (item 2.13 of the MDRS-UPDRS > 0 in usual life), 2) stability of others medical condition or that do not interfere with the research protocol and 3) effective contraception for woman of childbearing potential. Twenty-three healthy age-matched control subjects were also included in the GB-MOV study (**Supplementary Table 2**).

These studies were performed in accordance with the declaration of Helsinki and Good clinical practice guidelines. The local ethics committee approved the studies (N° 2012-A00225-38 and 2019-A01717-50), and all participants signed written informed consent to participate. These studies were registered on a clinical trial website (ClinicalTrials.gov: NCT01682668 and NCT04223427).

### Surgical procedure and anatomical localization of recording dipoles

Patients underwent a comprehensive assessment at the time of enrollment (baseline), followed by bilateral STN-DBS implantation. The two electrodes were implanted the same day, as previously reported, with direct targeting of the STN using 3D T2 Flair weighted images on preoperative 1.5 or 3Tesla MRI, and an additional indirect targeting using the basal ganglia atlas for patients operated at the Salpêtrière hospital^3^. Nineteen patients received quadripolar unsegmented electrodes with 4 contacts (electrode model 3389, Medtronic, Minneappolis, MN) and 19 received segmented electrodes with 8 contacts (electrode model Cartesia DB-2202, Boston Scientific, Marlborough, MA).The electrodes were connected to externalized cables to allow LFP recordings in the days following surgery (1-4 days, Fig. 1a-b), before implantation of the neurostimulator (Activa, Medtronic or Vercise PC, Boston Scientific). A 3D helical CT scan was performed the days following surgery to confirm the absence of early surgical complications and visualize electrode tracks.

**Fig. 1.**
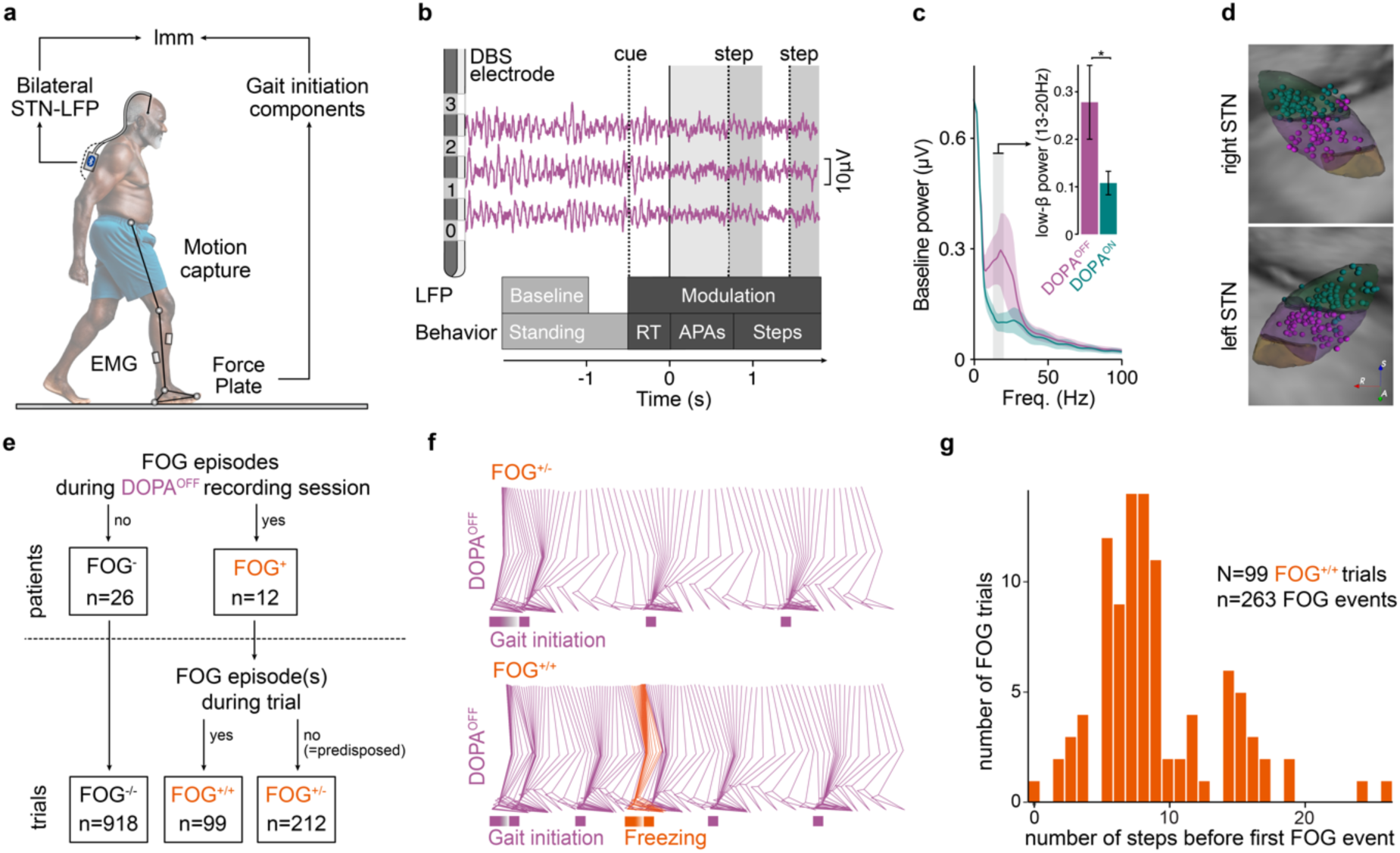
Recordings of subthalamic local field potentials with gait. **a.** Schematic representation of instrumented gait recordings using a force plate and an optical motion capture system with reflective markers positioned on all four limbs, together with STN LFP recording using an amplifier positioned in a small backpack. **b.** STN-LFP recorded through implanted DBS electrodes during a walking trial, aligned on gait initiation, with the cue, the delay (reaction time, RT) between the cue and the start of the anticipatory postural adjustment phase (APAs), and stepping. **c.** Average power spectra across DOPA^OFF^ and DOPA^ON^ conditions, at baseline, and the average low beta band power (inset). **d.** Location of the recording sites within the posterior-sensorimotor (green) and central-associative (purple) areas of the right and left STN. The different STN territories are represented in light green (posterior-motor), pink (central-associative) and yellow (anterior-limbic). **e.** Classification of the 38 patients depending on FOG occurrence during gait recordings in the DOPA^OFF^ condition. **f.** Reconstruction of leg kinematics in one individual FOG^+^ patient for one trial without FOG (FOG^+/−^, upper panel) and one with FOG occurring after gait initiation (FOG^+/+^, bottom panel, orange). **g.** Distribution of the onset time (after gait initiation) of the first FOG episode in trials with FOG. In FOG^+^ patients, we recorded 99 gait trials containing FOG episodes (FOG^+/+^ trials), in which 263 FOG episodes were detected.

The post-operative CT images were co-registered with the pre-surgical T1-weighted MRI to visualize the electrodes and contacts location within the STN area, using a detailed basal ganglia atlas for each individual patient. Recordings sites (dipoles) were identified as the location equidistant from the two contacts forming each bipolar montage. Recording sites were separated into those in the posterior-sensorimotor part of the STN and those in the central-associative part based on divisions inferred from external globus pallidus afferents to the STN.^3^ We then registered all the dipoles in the common brain space using a basal ganglia atlas that contains a visualization of the STN anatomo-functional territories^3^. For subsequent analyses, we retained only recording sites for which both contacts of the bipolar montage were localized within the STN (267 of 454), with 139 dipoles located in the dorsal-posterior-sensorimotor part of the STN and 128 dipoles located in a more ventral-anterior-central part of the STN (**Fig. 1d**).

### Experimental protocol

Participants underwent biomechanical and physiological recordings in an instrumented gait platform. Patients were assessed both without (DOPA^OFF^, after a 12-hour interruption of antiparkinsonian medication) and with dopamine medication (DOPA^ON^, after the administration of a suprathreshold dose of levodopa that corresponds to the usual morning levodopa dosage + 50 mg). In PD patients, the clinical severity of motor disability and gait and balance disorders were assessed using the MDS-UPDRS part 3, ‘axial’ (sum of the items: “speech”, “arising from chair”, “gait” “freezing of gait”, “postural stability” and “posture”), and Gait and Balance Scale (GABS) scores, both in the DOPA^OFF^ and DOPA^ON^ conditions (**Supplementary Table 1**).

### Gait recordings

Kinetics and kinematics parameters of gait were recorded 1 to 4 days after DBS surgery. Patients, barefoot, were instructed to commence walking on a force platform (0.9 × 1.8 m, AMTI, Advanced Mechanical Technology Inc. Watertown, MA, USA) and for 6-8 meters following a visual cue, make a half-turn, and return to their initial position (Supplementary Fig. 2a). Ground reaction forces and moments of gait initiation were recorded and sampled at 1000 Hz. Straight-ahead walking and turn were recorded using a motion capture system using 32 markers (Vicon®, Oxford, UK). Electromyographic activity (EMG) from the soleus and tibialis anterior from each leg were also recorded using surface electrodes and sampled at 2000 Hz. Patients performed on average 24 trials per treatment condition.

### STN LFP recordings

LFPs were obtained using bipolar montages formed from adjacent bipolar contact pairs. The Medtronic 3389 electrode has four contacts (1.27 mm diameter, 1.5 mm height, separated by 0.5 mm; the most ventral labelled 0 and the most dorsal 3) yielding 3 recording sites (0-1, 1-2, 2-3). The Cartesia DB-2202 lead has two ring contacts (contact 1 and 8, most ventral and dorsal, respectively) and two segmented contacts (2-3-4 and 5-6-7) allowing the recording of 9 dipoles (1.5mm² surface area, 1.3 mm diameter, 1.5 mm high, separated of 0.5 mm). The segmented contacts cover 90 degrees each and are separated by 30 degrees.

LFPs were amplified, filtered (band-pass filter: 0.25-300 Hz), sampled at 512 Hz, and transmitted via Bluetooth to the recording computer using an amplifier (Porti 32, TMS International, Enschede, The Netherlands) placed in a small backpack worn by the patient (**Fig. 1a**).

Gait kinetics and LFPs signals were simultaneously recorded and exported for off-line analyses using the MATLAB (MathWorks Inc.) toolbox (http://biomechanical-toolkit.github.io).

## Signal processing and analysis

### Gait parameters and FOG events

We focused our analysis of gait parameters on gait initiation, which has been shown to be a good proxy for studying the FOG state^35,36^ and allows to differentiate various components of gait and balance control^30^. We obtained 17 kinetic parameters from the force platform data using Nexus software (Version 10.2, NEXUS Software, Chesterland, OH, USA), which provided continuous signals proportional to the ground reaction forces (Rx, Ry, Rz in Newton) and moments (Mx, My, Mz in Newton meter) with respect to the mediolateral (x), anteroposterior (y) and vertical (z) axes of the force plate.^32^ Calling the subject’s mass m and the subject’s weight w, the mediolateral displacements of the centre of foot pressure (CoP) will then be defined as Mx/m and Center of mass (CoM) acceleration as (Rx-w)/m. For the APAs, we computed: 1) their duration, 2) the antero-posterior, 3) medio-lateral, and 4) vertical displacements of the CoP. At the time of foot off, we computed 5) the CoM forward speed. Following the gait initiation cycle, we then computed 6) the speed, 7) length, 8) width of the first step. We also introduced 5 timing parameters: the duration of 9) the first step, 10) double stance, and 11) second step, as well as 12) the cadence. As meaningful parameters of active balance control, we also computed the CoM downward speed 13) at foot strike (V1), and the corresponding 14) breaking of the fall compared to its maximum value during the step (V2). Finally, we computed during stepping 15) the maximum antero-posterior CoP speed and 16) its timing, and 17) the maximum medio-lateral displacement (**Supplementary Fig. 2b**). We used principal component analysis (PCA) using all trials to reduce dimensionality of these 17 kinetic parameters. Each varimax-rotated principal component (RPC score) was interpreted by exploring how scores across participants correlated with the 17 kinetic parameters using a Pearson’s correlation test (**Supplementary Fig. 2c**).

After exclusion of gait initiation trials for which we were not able to precisely detect the first biomechanical event (t0) corresponding to APAs onset (due to patients’ movements or false departure, or technical issues), we obtained 1229 gait trials in the DOPA^OFF^ and 1251 gait trials in the DOPA^ON^ conditions.

Kinematics recordings of straight-ahead walking and turn were analyzed to precisely determine the onset timing of FOG episodes, characterized by an arrest in walking flow and an inability to lift the foot from the ground. The occurrence of each FOG episode was also documented by the examiner during the recording sessions for every trial.

### Electrophysiological data processing

STN-LFPs were first high-pass filtered at 1 Hz and notch filtered to remove 50 Hz line noise, epoched into walking trials, and automatically and visually inspected for artifacts. Each trial was aligned from 2 seconds before the cue to the end of the second swing phase. Signals were transformed to the time-frequency domain using a multi-taper estimation algorithm implemented in the Chronux library in Matlab (http://chronux.org/). Power was calculated between 0 and 100 Hz using 3 orthogonal tapers using a 500 ms window stepped by 30 ms. Time-frequency maps were normalized by dividing all points by a baseline spectrum obtained by averaging the time-frequency map over a 800 ms period preceding the trial start. Power was transformed to decibels (10xlog10) for subsequent analyses.

All time-frequency maps obtained for each individual gait trial were visually inspected by at least two independent examiners (MLW, CO and MY) to exclude traces with obvious movement artifacts and/or saturated signals.

## Statistical analyses

### Gait performance in people with PD and effects of levodopa treatment

To assess the changes in gait RPCs scores across FOG groups (FOG^+^ vs. FOG^−^) and changes with dopaminergic treatment conditions (DOPA^OFF^ vs. DOPA^ON^), we used a two-way Anova with interaction and post-hoc paired t-tests across dopaminergic treatment conditions. To examine the link between gait RPCs scores, we used linear Pearson correlation tests for each FOG groups and treatment conditions.

To assess the changes in clinical scores (MDS-UPDRS part 3, axial and GABS scores) across FOG groups (FOG^+^ vs. FOG^−^) and changes between treatment conditions (DOPA^OFF^ vs. DOPA^ON^), we used a two-way ANOVA with interaction and post-hoc Wilcoxon rank-sum tests to identify differences between conditions.

### Correlation between gait and STN neuronal activity (LFP)

We examined the relationship between gait RPCs scores and STN LFPs using linear mixed models (lmm) that included spectral power as the dependent variable, 3 categorical predictors: 1) FOG^−^ vs. FOG^+^, 2) posterior vs. central STN subareas, 3) DOPA^OFF^ vs. DOPA^ON^, and one continuous predictor 4) the component score. We added an interaction term between the four predictors. The model included random intercepts with the recording dipole nested within subjects. When localization (posterior vs. central STN) within the STN was compared, a similar model was fit without the localization predictor. Models were fit using the lme4 package, and we estimated the slope between “RPC score” and “Spectral power” with the emtrends function of the emmeans package.

We ran the same analysis on the time course of frequency bands of interest (e.g., low beta: 13-20Hz). For each time point, we illustrate the estimated coefficient value (β) ± its standard error is reported, as well as the t-statistics (testing that the coefficient is equal to 0) along with its p-value.

We performed all statistical tests using R software 4.2.2. We corrected multiple comparisons to control the false discovery rate, and considered adjusted p-values < 0.05 as significant, except for time-frequency maps where we used a threshold at 0.001.

## Results

### Gait initiation performance relates to FOG predisposition and occurrence

During the DOPA^OFF^ recording session (n=1229 trials), 12 patients experienced FOG during straight-ahead walking or turning (FOG^+^ patients), while 26 patients did not (FOG^−^ patients). FOG^+^ patients demonstrated more severe gait and balance disorders (higher axial and GABS scores) compared to FOG^−^ patients (DOPA^OFF^), with no other significant demographic or clinical differences (**Supplementary Table S1, Supplementary Fig. 1**). Dopaminergic treatment significantly improved parkinsonian motor signs and gait and balance disorders in both groups to similar levels (**Supplementary Table 1, Supplementary Fig. 1**).

In the 12 FOG^+^ patients, 99 trials included FOG events (FOG^+/+^ trials, DOPA^OFF^ condition), with a median onset of 8 steps (IQR 6-11.5) after gait initiation. A total of 263 FOG episodes were recorded (**Fig 1e-g**). These FOG^+^ patients also completed 212 walking trials without FOG (FOG^+/−^ trials, **Fig 1e**). Among the 26 remaining FOG^−^ patients, 918 trials were recorded (FOG^−/−^ trials, DOPA^OFF^ condition, **Fig 1e**). During the DOPA^ON^ recording session, freezing was fully resolved in all FOG^+^ patients, except for one who experienced some residual FOG (patient M_0102 **Supplementary Table S1**), these trials were excluded from further analysis. In total, 1251 trials were recordedduring the DOPA^ON^ condition (FOG^+^ patients, n= 349; FOG^−^ patients, n= 902).

Principal component analysis (PCA) with varimax rotation of gait parameters from trials without FOG (n=1130 trials) revealed five orthogonally RPC accounting for 73% of variance (**Fig. 2**, **Supplementary Fig. 2**). Scores for RPCs 1, 2 and 5 were significantly modified by dopaminergic medication (DOPA^ON^-DOPA^OFF^ estimate [SE] for RPC1=0.72 [0.11], p<1.17e-07; RPC2=0.55 [0.14], p=4.64e-04; RPC5=-0.62 [0.12], p=1.35e-05), while scores for RPCs 3 and 4 showed no significant changes (RPC3=0.039 [0.14], p=0.813; RPC4=0.31 [0.17], p=0.322, **Supplementary Fig. 2**). RPC1 (24% of total variance) captured swing speed and other variables related to forward stepping vigor and propulsion, which we labelled *pace* (**Fig 2a**). RPC2 (19% of total variance) reflected step times and cadence, which we labelled *rhythm*. RPC5 (7.5% of total variance) represented step width and lateral center-of-mass speed, indicating lateral balance control or body sway, which we labelled *balance*. Comparisons with healthy age-matched controls indicate that dopaminergic treatment improved *pace* and *rhythm* but worsened *balance* (**Fig 2b-c, Supplementary Fig. 2**).

**Fig. 2.**
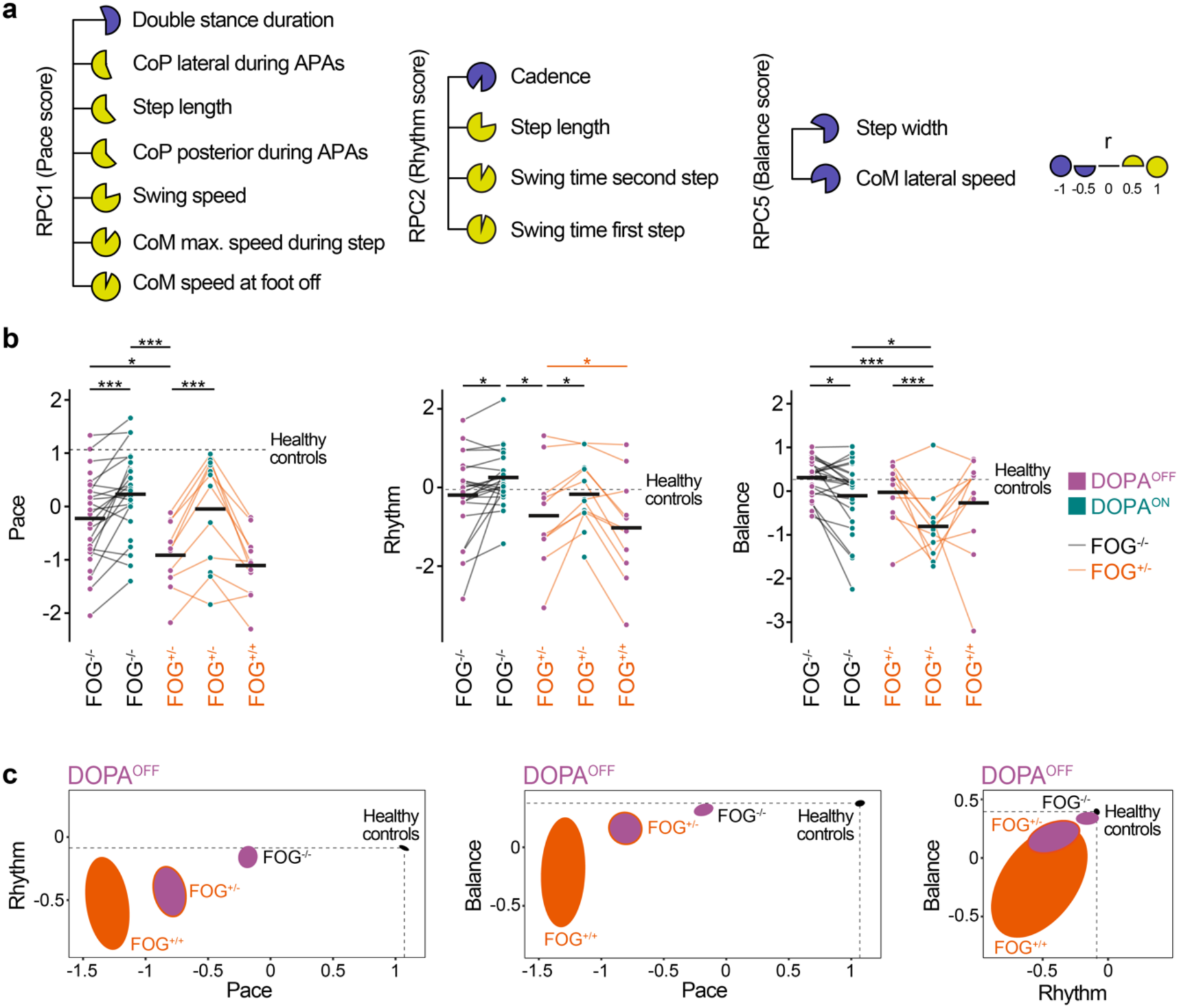
Decomposition of gait initiation using principal components. **a.** Results of the principal component analysis with correlation (r scores) for each RPC. RPC1 strongly correlated with speed and step length, thus reflecting walking pace, RPC2 correlated with cadence and step times, reflecting stepping rhythm, and RPC3 correlated with step width and lateral speed, reflecting balance. **b.** Scores for pace (RPC1), rhythm (RPC2) and balance (RPC3) obtained in the DOPA^OFF^ (purple) and DOPA^ON^ (green) condition for patients without FOG (FOG^−/−^ trials) and for patients who experienced FOG (FOG^+^), separating trials without (FOG^+/−^ trials) and with FOG (FOG^+/+^ trials). Each dot represents one individual PD subject and the black lines the means. The dashed horizontal line shows the mean score obtained in healthy aged-matched controls. \**P*<0.05, \*\**P*<0.01, \*\*\**P*<0.001. **c.** Correlation analysis between RPC scores in the DOPA^OFF^ condition. Ellipses show mean ± s.d. of correlation scores projected onto pairs of RPCs. The black ellipse is the correlation score obtained in healthy aged-matched controls.

In the DOPA^OFF^ condition, FOG^+^ patients exhibited significantly lower scores for *pace* (RPC1) than FOG^−^ patients (FOG^−/−^ trials-FOG^+/−^ trials estimate [SE]=0.75 [0.30], p=0.025), **Fig 2b**). *Rhythm* (RPC2) and *balance* (RPC5) scores showed no significant differences (RPC2 =0.70 [0.40], p=0.138; RPC5=0.25 [0.19], p=0.406). Within FOG^+^ patients, we found significantly lower scores for *rhythm* in FOG^+/+^ trials compared to FOG^+/−^ trials (RPC2, estimate [SE]=-0.243 [0.102], p=0.035; **Fig. 2b**), indicating degraded stepping rhythm at gait initiation, with no significant differences in *pace* or *balance* scores (estimate [SE] RPC1=-0.102 [0.089], p=0.246; RPC5=-0.099 [0.179], p=0.585). Dopaminergic medication improved *pace* (RPC1) and *rhythm* (RPC2) scores in both groups, eliminating significant differences between FOG^+^ and FOG^−^ patients (DOPA^ON^ condition, FOG^−^ – FOG^+^ estimate [SE] for RPC1=0.30 [0.30], p=0.391; RPC2=0.42 [0.26], p=0.14, **Fig 2b**). However, *balance* scores (RPC5) worsened in both groups under dopaminergic treatment (DOPA^ON^ – DOPA^OFF^ estimate [SE] for FOG^−^ patients=0.39 [0.13], p=0.013; FOG^+^ patients: estimate [SE]=0.85 [0.21], p=6.16e-04), with a significantly greater worsening in FOG^+^ patients compared to FOG^−^ patients (RPC5 estimate [SE]=0.71 [0.28], p=0.022, **Fig 2b**). We found no significant differences in RPC3 and RPC4 scores among patient groups and across dopaminergic treatment condition (**Supplementary Fig. 2**).

To investigate the differences and behavioral strategies for successful gait initiation between FOG^−^ and FOG^+^ patients in the DOPA^OFF^ condition, we explored the link between *pace*, *rhythm* and *balance* scores (**Fig 2c**). *Pace* (RPC1) and *balance* (RPC5) scores were positively correlated in FOG^−^ patients (r=0.27, t_(916)_=8.44, p_c_<0.001), but this correlation was absent in FOG^+^ patients (r=-0.04, t_(210)_=0.61, p_c_=0.54). Instead, FOG^+^ patients displayed a significant negative correlation between *rhythm* and *pace* (r=-0.22, t_(210)_=3.19, p_c_=0.0022) and a significant positive correlation between *rhythm* and *balance* (r=0.38, t_(210)_=6.06, p_c_<0.001). These relationships degraded further in FOG^+/+^ trials (**Fig. 2c**).

### Subthalamic activity correlates with FOG predisposition and is spatially distributed

We examined the relationship between LFP spectral power and gait RPC scores, focusing first on trials without FOG in the DOPA^OFF^ condition. The analysis included data from dipoles located within the STN (n=267 dipoles), comprising 946 traces (725 in FOG^−^ and 221 in FOG^+^ patients, **Fig. 1d**, **Fig. 3**).

**Fig. 3.**
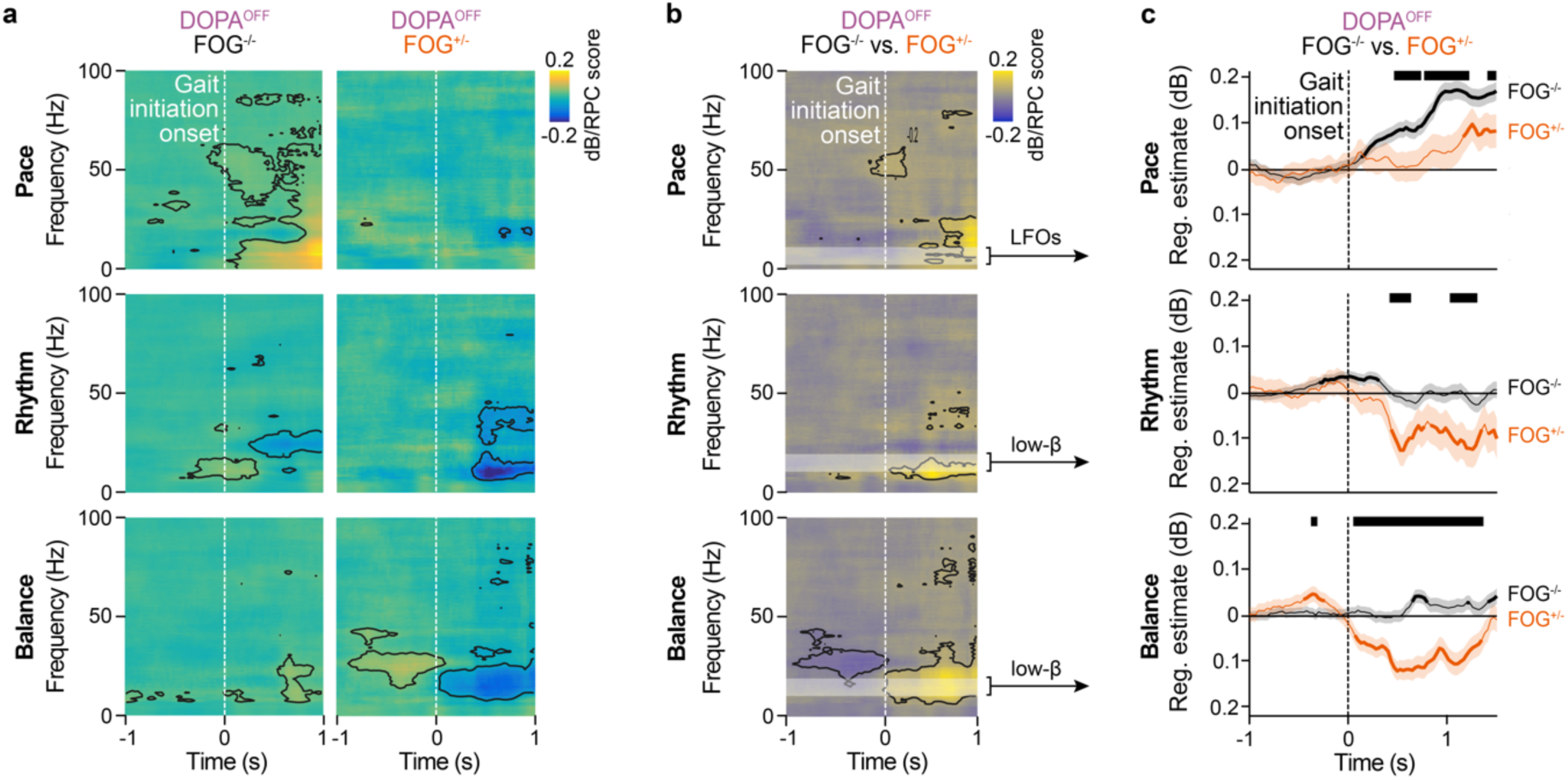
Relationship between subthalamic neuronal activity and gait initiation parameters. **a.** Time-frequency maps illustrating associations between STN LFP and pace (RPC1), rhythm (RPC2) and balance (RPC5) scores in the DOPA^OFF^ condition, for both FOG^−^ patients (FOG^−/−^ trials, n=658) and FOG^+^ patients in the absence of FOG episodes (FOG^+/−^ trials, n=233). The white line indicates the start of the APAs phase. Black contours outline regions of significance. **b.** Comparison between FOG^−/−^ and FOG^+/−^ trials (difference in slopes). **c.** Mean associations between low frequency oscillations (theta-alpha [4-12 Hz] bands) and pace (RPC1, upper panel), for FOG^−/−^ trials (FOG^−^ patients, black trace) and FOG^+/−^ trials (FOG^+^ patients, orange trace). The black bars illustrate significant differences between the FOG^−/−^ and FOG^+/−^ trials. Middle and bottom graphs report the same analysis for low beta band (13-20 Hz) power associated with rhythm (RPC2) and balance (RPC5) scores.

We observed transient positive associations between gait scores and theta/alpha (4-12 Hz) and gamma bands (>35Hz) power, and transient negative associations with beta band power (13-35Hz; **Fig 3a, Supplementary Fig. 3**). Comparing the two patients groups, we found that FOG^+^ patients exhibited weaker positive associations between theta/alpha and gamma band powers and *pace* scores and stronger negative associations between low beta band power and *rhythm* or *balance* scores following gait initiation, which were absent or weaker in FOG^−^ patients (**Fig. 3b-c**). This negative beta band associations extended into the high beta band, distinguishing FOG^+^ from FOG^−^ patients (**Fig. 3a-b**).

Examining LFP modulations and associations across posterior-sensorimotor and central-associative STN subareas (**Fig 1d, Supplementary Fig. 4**) in the two patients groups, we found that FOG^+^ patients exhibited: 1) stronger negative associations between low beta power and *pace* score in the posterior STN (**Fig. 4a-c, Supplementary Fig. 5**); 2) negative associations between low beta power and *rhythm* in both the posterior and central STN subareas, contrasting with positive associations in the central STN for FOG^−^ patients (**Fig 4d-f, Supplementary Fig. 5**); and 3) negative associations between low beta power and *balance* score in both STN subareas, with stronger effects in the posterior STN, differing from positive associations observed in FOG^−^ patients (**Fig. 4g-i**, **Supplementary Fig. 5**). FOG^+^ patients were further distinguished by a pronounced positive correlation between beta band activity and *balance* score starting before gait initiation, which was significantly larger in the posterior STN (**Fig. 4g-i**, **Supplementary Fig. 5**).

**Fig. 4.**
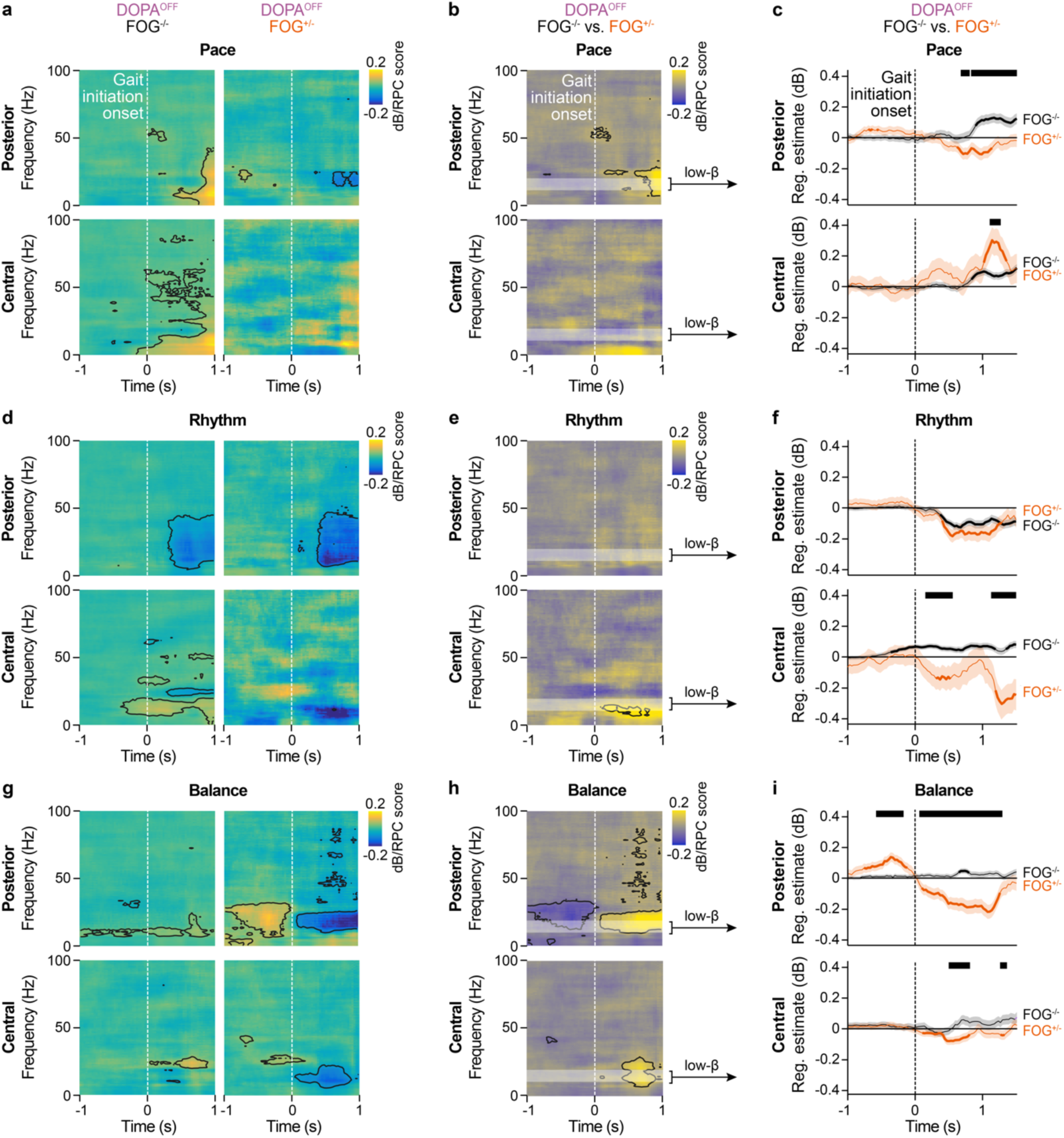
Low beta modulation during gait initiation is spatially distributed within the STN and differently affected between FOG^−^ and FOG^+^ patients. **a.** Time-frequency maps illustrating associations between STN LFP recorded within the posterior and central STN and pace (RPC1) in the DOPA^OFF^ condition, in patients without FOG^−^ (FOG^−/−^ trials) and FOG^+^ patients in the absence of FOG episodes during trials (FOG^+/−^ trials). **b.** Comparison between patient groups (difference in slopes). **c.** Mean associations between low beta band (13-20 Hz) and stepping rhythm (RPC2) for FOG^−/−^ trials (FOG^−^ patients, black line) and FOG^+/−^ trials (FOG^+^ patients, orange line). **d-f,** Same analysis as in **a-c**, for associations between STN LFP and rhythm (RPC2) scores, and **g-i** for associations between STN LFP and balance (RPC5) scores.

### Predictive patterns of FOG occurence

To explore STN dysfunction linked to FOG, we compared trials with freezing (FOG^+/+^) to those without (FOG^+/−^) among FOG^+^ patients (DOPA^OFF^ condition, **Fig 5**). We observed stronger negative associations between high beta band power and *pace* score in FOG^+/+^ vs. FOG^+/−^ trials, significant only in the posterior STN (**Fig 5a-c, Supplementary Fig. 6**). In contrast, positive association between *rhythm* and both high beta and theta band activity emerged in the posterior STN during FOG^+/+^ trials, but were absent or reversed in FOG^+/−^ trials (**Fig 5d-f**). For *balance*, we found a stronger negative association between high beta band power in the central STN and positive association with gamma band power in the posterior STN (**Fig 5g-i**).

**Fig. 5.**
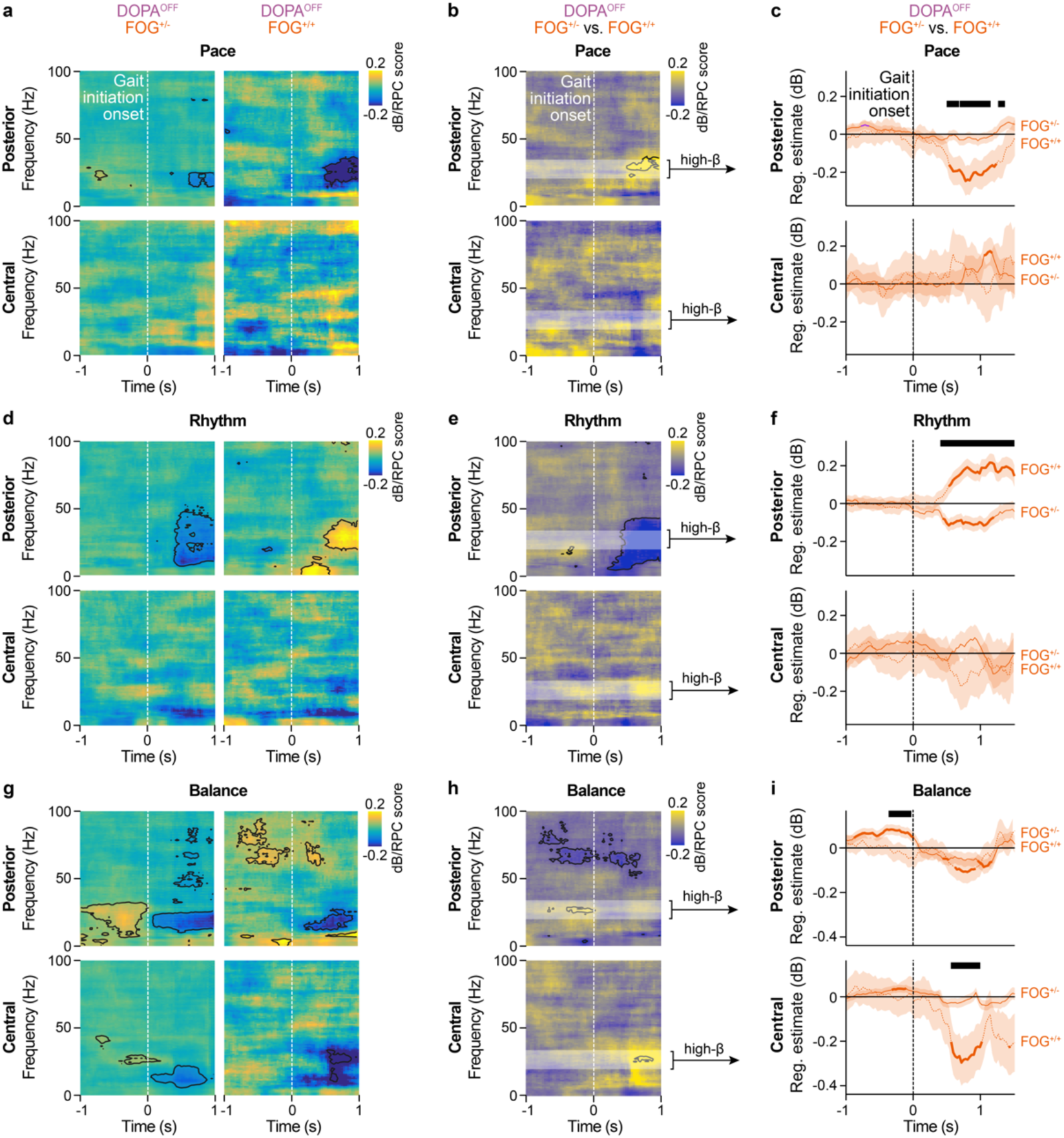
Subthalamic signature at gait initiation correlates with eventual FOG occurence. **a-c,** Same analysis as for Fig 4 but comparing between FOG^+/−^ and FOG^+/+^ trials in FOG^+^ patients for the pace score, **d-f,** for rhythm and **g-i** for balance. Legends are the same as for Fig. 3 and 4.

### Dopaminergic medication improves gait disability and reestablishes STN activity

In the DOPA^ON^ condition, we analyzed 875 traces (633 from FOG^−^ patients, 242 from FOG^+^ patients). We found positive associations between low beta band power and *pace* score in FOG^−^ patients, with minimal changes between DOPA^OFF^ and DOPA^ON^ condition (**Fig 6a-b**). In FOG^+^ patients, dopamine treatment shifted associations between low beta band power and *pace* from negatice (DOPA^OFF^) toward positive, aligning more closely with FOG^−^ patients patterns (**Fig 6c-d**). For *rhythm*, the negative association with beta band power observed in FOG^+^ patients was reduced with dopamine treatment (**Fig 6c-d)**. For *balance*, dopaminergic treatment mitigated the negative association between low beta power observed in the DOPA^OFF^ condition in FOG^+^ patients, resembling associations seen in FOG^−^ patients in the DOPA^ON^ condition (**Fig. 6a-b**).

**Fig. 6.**
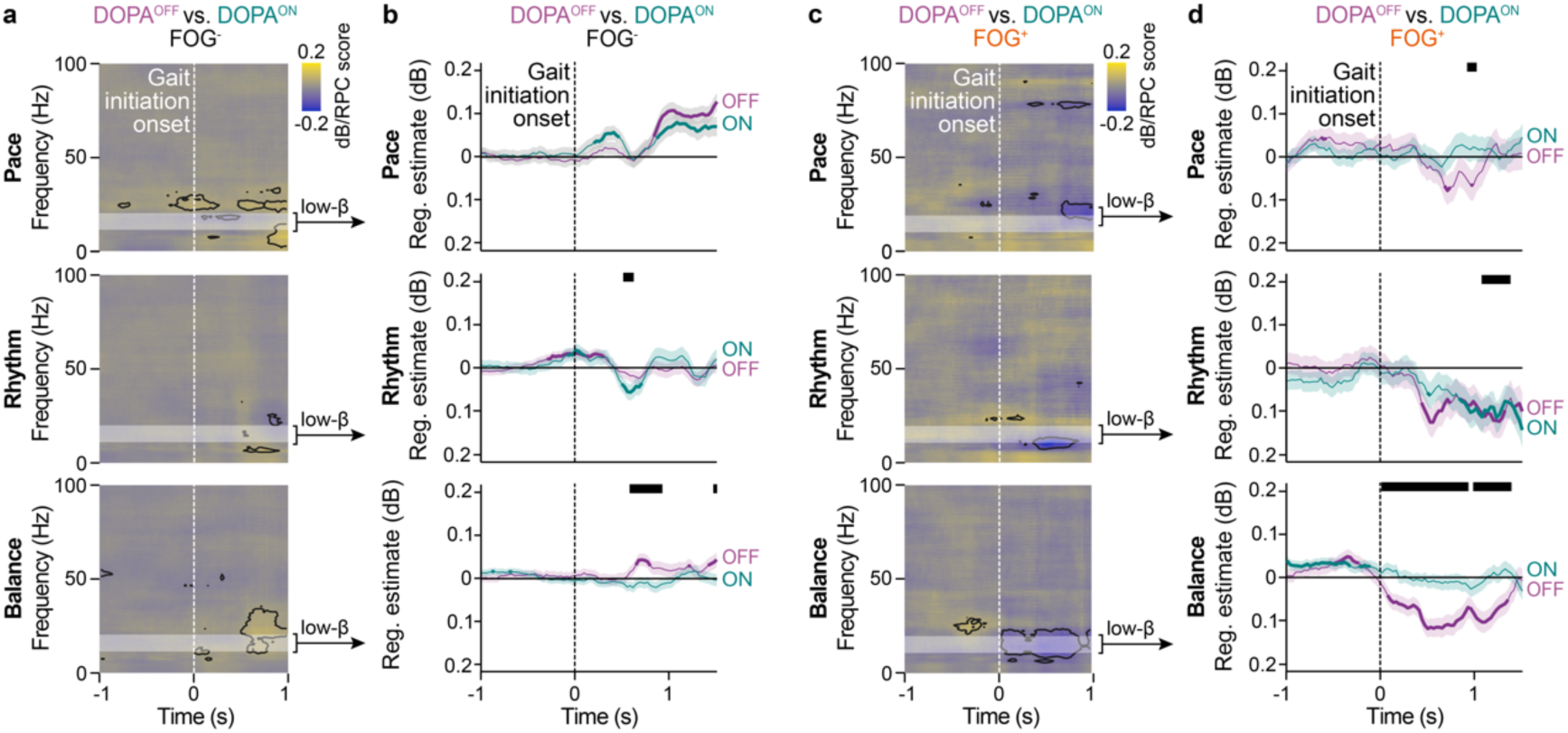
Effects of dopaminergic medication on the subthalamic neuronal activity at gait initiation. **a.** Same analysis as Fig 3 for the DOPA^ON^ condition, and the contrast between DOPA^ON^ and DOPA^OFF^ conditions in both **b and c,** FOG^−^ and **d and e**, FOG^+^ patients. Legends as for Fig. 3.

## Discussion

This study distinguishes between two FOG states—predisposition and occurrence—each characterized by distinct gait initiation strategies and STN neuronal activity patterns. FOG^+^ patients utilize a compensatory strategy, relying on steady stepping *rhythm* modulated by low beta activity within posterior-sensorimotor and central-associative STN subareas to initiate and sustain gait. When this stepping rhythm strategy deteriorates further, FOG episodes occur and is marked by strong associations with high beta power in the posterior-sensorimotor STN and diminished low beta STN negative associations across the posterior and central STN. Dopamine treatment improved gait patterns and partially restored STN neuronal activity, reducing the differences between FOG^+^ and FOG^−^ patients.

### Behavioral encoding of FOG predisposition correlates with STN neuronal activity

Differences in gait initiation patterns have been previously demonstrated in PD FOG^+^ patients and linked to dysfunction in medial pre-motor cortical areas, particularly within the SMA. This dysfunction impairs postural adjustments during gait initiation^37^, with reduced stepping pace and gait automaticity^38,39^. In FOG^+^ patients, we observed weaker associations between theta-alpha power associations with stepping pace. In healthy people, alpha rhythms at the cortical level appear to inhibit neuronal populations that could interfere with movement selection ^40^. In PD patients, cortical alpha desynchronization occurs during walking^41^, with alpha synchronization in the pedunculopontine nucleus correlating with walking speed and inversely with gait impairment and FOG^42^. Alpha band synchronization within the STN has also been reported in PD^7^, associated with movement initiation^43^, and found coupled with the temporal and parietal cortices, brainstem, and cerebellum^44,45^. Synchronized STN-cortical theta-alpha band power was reported during walking,^25,27^ particularly before and during contralateral leg swing^46^. This synchronization, preferentially distributed in the anterior-medial STN^7,47^, is also linked to non-motor cognitive and emotional processing^47,48^. These observations suggest that in FOG^−^ patients, the correlation between stepping pace and low frequency STN activity may reflects increased attention and resource allocation during gait initiation^49^, and that the reduced pace and lower STN alpha power modulations observed in FOG^+^ patients would also reflect additional cortico-STN-MLR pathway dysfunction.^45,50,51^

FOG^+^ patients exhibited more extensive and low beta desynchronization related to rhythm and balance at gait initiation. Low beta activity has been identified as a key frequency band in parkinsonian motor signs in the DOPA^OFF^ state, exhibiting dynamic changes across both physiological and pathological motor states^9,12^. In animals, modulating STN neural activity alters locomotor activity^52,53^, and abolishing excessive STN beta synchronisation in parkinsonian mice ameliorates locomotion^54,55^. In PD, STN beta band power decreases during gait^23,46^, and encodes volitional commands for leg muscles activation and vigor^21^. This indicates that FOG^+^ patients likely compensate for deficits in gait rhythmicity, transition and balance^38,56^, by amplifying the negative association between low beta band power and gait initiation, enhancing stepping rhythm and employing volitional gait encoding strategies^21^. This compensatory activity was observed across both posterior-sensorimotor and central-associative subareas, whereas low beta band modulation was restricted to the posterior STN in FOG^−^ patients. The borader spatial distribution of low beta modulation in FOG+ patients likely reflects increased cortical engagement, recruiting additional cortical (i.e. prefrontal) areas for voluntary gait control. Recent studies have linked low beta oscillations in the medial cortex, including the SMA, which controls gait initiation and postural adjustments^57^. This activity, primarily subcortical ^57^, reflects basal ganglia dysfunction from dopaminergic loss^8,9^. Consistent with this, dopaminergic therapy (DOPA^ON^ state) significantly improved motor motor performance^8,9,58^, including stepping rhythm and walking pace, and reduced low beta power associations in both patient groups.

In FOG^+^ patients, low beta desynchronization was also associated with balance control, which was more impaired compared to FOG^−^ patients and worsened in the DOPA^ON^ state^59,60^. Balance deficits in PD are tied to disruptions in attentional control and sensory or visual-vestibular information processing^61,62^, potentially related to additional cholinergic lesions^61,63^. Leg proprioceptive integration was recently linked to beta band modulation in the STN^21^, and modulation of sensory feedback using lumbosacral electric epidural stimulation in one PD patient with FOG have shown encouraging results^64^.

### FOG signature within the STN

During walking trials where FOG^+^ patients experienced FOG episodes, profound alterations in stepping rhythm at gait initiation were observed, including increased cadence and shorter walking durations. Previous studies have documented similar degradations in stepping rhythm during straightforward gait in FOG patients, such as increased cadence, loss of lower limb alternation, and asymmetrical step length^38^. Additionally, altered gait patterns often precede FOG onset during experimental tasks like obstacle avoidance or door passage^65,66^. In our patients, FOG episodes were preceded by an increased association between high beta band activity and stepping rhythm in the posterior part of the STN, coupled with an inverse association between high beta band activity and balance in the central STN. These findings suggests that disruptions in STN activity, particularly within the high beta band, arise during initiation of walking and are predictive of FOG occurence. Previous research has linked STN beta oscillations to FOG, reporting increased entropy^19^, elevated low or high beta power^18,21,25,26^, prolonged STN beta burst durations^10^, and cortico-STN decoupling^27^. High beta-gamma phase-amplitude coupling in the primary motor cortex has also been associated with FOG episodes^67^. Our recordings also showed that FOG episodes were marked by elevated high beta power associations in the posterior STN – primarily connected with motor and premotor cortical areas^68^ – well before the onset of a FOG episode. Concurrently, a significant decrease in high beta correlation with balance in the central STN suggests additional functional disconnection between the posterior and central STN subareas, which could precipitate FOG.

Subthalamic high beta power coupling with medial sensorimotor cortical areas has been reported in PD patients, with cortical activity leading the STN^44^. This high beta activity is not influenced by dopaminergic medication^44,57^ and is though to induce increased surround inhibition within the BG pathways. This inhibition would block movement facilitation typically promoted by the direct pathway, which is weakened by dopamine depletion in PD^69–71^. The hyperdirect pathway plays a key role when multiple competing motor programs are active in the premotor cortex^72^, globally inhibiting prepared responses as behavioral demands arise^73^. The differential changes in low and high beta associations with gait scores observed in our FOG^+^ patients may reflect a physiological conflict within the STN. This confkict, arising between competing motor programs^70,72^, likely results in over-inhibition of the “center-facilitated” gait initiation program^69^, ultimately triggering FOG episodes.

### Gait pattern and STN neuronal modulations: potential biomarkers for adaptive deep brain stimulation to treat FOG

In FOG^+^ patients, both gait patterns and STN modulations associated with FOG emerge during gait initiation, well before the onset of FOG episodes. This observation suggests that external triggers (e.g., gait patterns) and internal cues (e.g. neuronal activity) could be harnessed to adapt STN-DBS settings proactively, preventing FOG episodes at the onset of walking. Recent clinical advances highlight the feasibility of closed-loop DBS, which adapts stimulation parameters based on behavioral outcomes or neuronal activity. This personalized approach adjusts DBS settings according to the patient’s state, potentially reducing side-effects and enhancing efficacy by minimizing unncecessary stimulation^74^. For example, in patients with essential tremor, wearable accelerometers have been used to detect postural tremor^75^ and control thalamic DBS, demonstrating comparable efficacy to continuous DBS^76^. Similarly, studies on thalamic and combined thalamic-cortical sensing in ET patients report superior outcomes compared to continuous DBS^77–79^. In PD patients, adaptive STN-DBS informed by kinetics parameters,^80^ beta-band power, or beta burst duration within the STN has shown benefits in pilot studies, improving upper limb akinesia, rigidity, tremor and reaching movements^33,81,82^, with benefits correlating to beta band suppression^82^. However, such adaptive approaches often require the development of optimized personalized classifiers^83^ to improve the responsiveness of closed-loop algorithms to normal and abnormal motor behaviors^84^. In this study, the limited number of recorded FOG episodes per patient presented challenges for developing individualized classifiers. To address this, future studies could induce additional FOG episodes in controlled gait lab conditions^21,27,85^ or capture more episodes under naturalistic settings^86^. Advances in sensing neurostimulators may further facilitate real-time adaptive DBS, enabling at-home implementation^87,88^.

### Study limitations

This study has several limitations. First, gait initiation and FOG episodes were recorded under controlled experimental conditions rather than in real-life settings. While this controlled approach ensures data consistency, it may not fully capture the variability and complexity of FOG episodes as they occur in daily life, potentially limiting the generalizability of the findings. Nonetheless to our knowledge, this study is the first to report such a large number of FOG episodes combining STN-LFP recordings and instrumented gait assessments. Second, this study did not include long-term follow-up data to evaluate the durability of STN modulation changes over time and their correlation with gait benefits of STN-DBS. This limits our understanding of the long-term effects of STN-DBS on gait and FOG. Future studies could address this by using advanced neurostimulators capable of continuously recording STN neuronal activity while applying effective STN-DBS. Additionally, we used a 3D histological and deformable atlas of the basal ganglia, registered to individual preoperative MRIs, to localize STN dipoles within STN subareas. While this method facilitated comparisons of anatomical data across patients by creating a common brain space, it may introduce some inaccuracies, particularly in patients with larger third ventricles, where electrode locations might vary slightly between individuals. However, we carefully verified electrode and dipoles location for each patient and included only those situated within the STN in the final analysis.

## Conclusion

Our results support the existence of a predisposed state for FOG in which patients experience a loss of gait automaticity. This loss appears to be compensated by increased volitional control on walking, leading to a more rhythmic stepping pattern to manage imbalance and facilitate foot landing. This compensation is linked to more broadly distributed low beta modulations within the STN. Ultimately, our results confirm the role of the STN in gait encoding, highlighting the importance of developing decoders tailored to transient states in PD patients. Specifically, modulating low beta activity through adaptative DBS in the central STN shows promise for improving walking in FOG^+^ patients. Furthemore, targeting high beta activity in the posterior STN could offer a more effective strategy for preventing FOG episodes. Recent adavances in adaptive DBS, coupled with improved data collection and refined algorithms, highlight the potential of leveraging both external gait patterns and internal neuronal activity as dynamic triggers, offering new hope for managing gait disorder and FOG for individual PD patients.

## Abbreviations

DBS: deep brain stimulation
DOPA: dopamine medication
FOG: freezing of gait
GABS: Gait and Balance Scale
LFP: local field potentials
MDS-UPDRS: Movement Disorders Society-Unified Parkinson’s disease Rating Scale
MLR: mesencephalic locomotor region
PD: Parkinson’s disease
RPC: Rotated Principal Component
STN: sutbhalamic nucleus

## Acknowledgements

The authors express sincere gratitude for the dedication demonstrated by our patients in participating in this research. We are also grateful to Anais Hervé and Sabine Meunier, and nurses of the Clinical Investigation Center at the Paris Brain Institute for their assistance in organizing the research program, to Xavier Drevelle, Dorian Banier, Willy Bertucchi, and Edward Soundaravelou for their help in data recordings, Yannick Mullié for his help in gait data analysis, and to David Maltête, Elodie Hainque and Julie Bourilhon for their help in patients’ recruitment.

## Authors contributions

M.L.W, A.V.H, C.K, B.L. designed the research; A.C.C, M.Y, H.B, A.V.H, A.D, C.O, D.Z, S.D, C.K., and M.L.W. acquired the data; A.C.C, M.Y, A.V.H, A.D, C.O, D.Z, S.F-V, K.L., B.L., and M.L.W. made the analysis and interpreted the data; A.C.C, M.Y, and M.L.W. made the first draft of manuscript; A.C.C, M.Y, C.K., B.L. and M.L.W. performed revision of the manuscript.

## Supplementary material

Supplementary material is available.

## References

1. Nutt, J. G. et al. Freezing of gait: moving forward on a mysterious clinical phenomenon. Lancet Neurol 10, 734–744 (2011).

2. Mirelman, A. et al. Gait impairments in Parkinson’s disease. Lancet Neurol 18, 697– 708 (2019).

3. Karachi, C. et al. Clinical and anatomical predictors for freezing of gait and falls after subthalamic deep brain stimulation in Parkinson’s disease patients. Parkinsonism Relat Disord 62, 91–97 (2019).

4. Fasano, A., Aquino, C. C., Krauss, J. K., Honey, C. R. & Bloem, B. R. Axial disability and deep brain stimulation in patients with Parkinson disease. Nat Rev Neurol 11, 98–110 (2015).

5. Hammond, C., Bergman, H. & Brown, P. Pathological synchronization in Parkinson’s disease: networks, models and treatments. Trends Neurosci 30, 357–364 (2007).

6. Oswal, A. et al. Deep brain stimulation modulates synchrony within spatially and spectrally distinct resting state networks in Parkinson’s disease. Brain 139, 1482–1496 (2016).

7. Darcy, N. et al. Spectral and spatial distribution of subthalamic beta peak activity in Parkinson’s disease patients. Exp Neurol 356, 114150 (2022).

8. Tinkhauser, G. et al. Beta burst dynamics in Parkinson’s disease OFF and ON dopaminergic medication. Brain 140, 2968–2981 (2017).

9. Mathiopoulou, V. et al. Modulation of subthalamic beta oscillations by movement, dopamine, and deep brain stimulation in Parkinson’s disease. NPJ Parkinsons Dis 10, 77 (2024).

10. Anidi, C. et al. Neuromodulation targets pathological not physiological beta bursts during gait in Parkinson’s disease. Neurobiology of Disease (2018) doi:10.1016/j.nbd.2018.09.004.

11. Morelli, N. & Summers, R. L. S. Association of subthalamic beta frequency sub-bands to symptom severity in patients with Parkinson’s disease: A systematic review. Parkinsonism Relat Disord 110, 105364 (2023).

12. Lofredi, R. et al. Subthalamic beta bursts correlate with dopamine-dependent motor symptoms in 106 Parkinson’s patients. NPJ Parkinsons Dis 9, 2 (2023).

13. Fischer, P. et al. Alternating Modulation of Subthalamic Nucleus Beta Oscillations during Stepping. J Neurosci 38, 5111–5121 (2018).

14. Tan, H. et al. Decoding Movement States in Stepping Cycles Based on Subthalamic LFPs in Parkinsonian Patients. Conf Proc IEEE Eng Med Biol Soc 2018, 1384–1387 (2018).

15. Toledo, J. B. et al. High beta activity in the subthalamic nucleus and freezing of gait in Parkinson’s disease. Neurobiol Dis 64, 60–65 (2014).

16. Georgiades, M. J. et al. Hitting the brakes: Pathological subthalamic nucleus activity in Parkinson’s disease gait freezing. Brain (2019) doi:10.1093/brain/awz325.

17. Welter, M.-L. et al. Virtual walking through a doorway promotes a beta-gamma power imbalance in the subthalamic nucleus in Parkinson’s disease. Clin Neurophysiol 162, 28–30 (2024).

18. Singh, A. et al. Freezing of gait-related oscillatory activity in the human subthalamic nucleus. Basal Ganglia 3, 25–32 (2013).

19. Syrkin-Nikolau, J. et al. Subthalamic neural entropy is a feature of freezing of gait in freely moving people with Parkinson’s disease. Neurobiol Dis 108, 288–297 (2017).

20. Arnulfo, G. et al. Phase matters: A role for the subthalamic network during gait. PLoS ONE (2018) doi:10.1371/journal.pone.0198691.

21. Thenaisie, Y. et al. Principles of gait encoding in the subthalamic nucleus of people with Parkinson’s disease. Sci Transl Med 14, eabo1800 (2022).

22. Canessa, A., Palmisano, C., Isaias, I. U. & Mazzoni, A. Gait-related frequency modulation of beta oscillatory activity in the subthalamic nucleus of parkinsonian patients. Brain Stimul 13, 1743–1752 (2020).

23. Hell, F., Plate, A., Mehrkens, J. H. & Bötzel, K. Subthalamic oscillatory activity and connectivity during gait in Parkinson’s disease. Neuroimage Clin 19, 396–405 (2018).

24. Storzer, L. et al. Bicycling suppresses abnormal beta synchrony in the Parkinsonian basal ganglia. Ann Neurol 82, 592–601 (2017).

25. Klocke, P. et al. Supraspinal contributions to defective antagonistic inhibition and freezing of gait in Parkinson’s disease. Brain 147, 4056–4071 (2024).

26. Chen, C. C. et al. Subthalamic nucleus oscillations correlate with vulnerability to freezing of gait in patients with Parkinson’s disease. Neurobiology of Disease (2019) doi:10.1016/j.nbd.2019.104605.

27. Pozzi, N. G. et al. Freezing of gait in Parkinson’s disease reflects a sudden derangement of locomotor network dynamics. Brain (2019) doi:10.1093/brain/awz141.

28. Zaidel, A., Spivak, A., Grieb, B., Bergman, H. & Israel, Z. Subthalamic span of beta oscillations predicts deep brain stimulation efficacy for patients with Parkinson’s disease. Brain 133, 2007–2021 (2010).

29. Telkes, I. et al. Local field potentials of subthalamic nucleus contain electrophysiological footprints of motor subtypes of Parkinson’s disease. Proc Natl Acad Sci U S A 115, E8567–E8576 (2018).

30. Cherif, S. et al. Directional subthalamic deep brain stimulation better improves gait and balance disorders in Parkinson’s disease patients: a randomized controlled study. Annals of Neurology (2024).

31. Fan, H. et al. Optimal subthalamic stimulation sites and related networks for freezing of gait in Parkinson’s disease. Brain Commun 5, fcad238 (2023).

32. Temiz, G. et al. Freezing of gait depends on cortico-subthalamic network recruitment following STN-DBS in PD patients. Parkinsonism Relat Disord 104, 49–57 (2022).

33. Little, S. et al. Bilateral adaptive deep brain stimulation is effective in Parkinson’s disease. J Neurol Neurosurg Psychiatry 87, 717–721 (2016).

34. Tinkhauser, G. et al. The modulatory effect of adaptive deep brain stimulation on beta bursts in Parkinson’s disease. Brain 140, 1053–1067 (2017).

35. Delval, A., Tard, C. & Defebvre, L. Why we should study gait initiation in Parkinson’s disease. Neurophysiol Clin 44, 69–76 (2014).

36. Schlenstedt, C. et al. Are Hypometric Anticipatory Postural Adjustments Contributing to Freezing of Gait in Parkinson’s Disease? Front Aging Neurosci 10, 36 (2018).

37. Jacobs, J. V., Lou, J. S., Kraakevik, J. A. & Horak, F. B. The supplementary motor area contributes to the timing of the anticipatory postural adjustment during step initiation in participants with and without Parkinson’s disease. Neuroscience 164, 877–885 (2009).

38. Plotnik, M., Giladi, N. & Hausdorff, J. M. Bilateral coordination of walking and freezing of gait in Parkinson’s disease. Eur J Neurosci 27, 1999–2006 (2008).

39. Yogev, G. et al. Dual tasking, gait rhythmicity, and Parkinson’s disease: which aspects of gait are attention demanding? Eur J Neurosci 22, 1248–1256 (2005).

40. Brinkman, L. et al. Independent Causal Contributions of Alpha– and Beta-Band Oscillations during Movement Selection. J Neurosci 36, 8726–8733 (2016).

41. Storzer, L. et al. Bicycling and Walking are Associated with Different Cortical Oscillatory Dynamics. Front Hum Neurosci 10, 61 (2016).

42. Thevathasan, W. et al. Alpha oscillations in the pedunculopontine nucleus correlate with gait performance in parkinsonism. Brain 135, 148–160 (2012).

43. Klostermann, F. et al. Task-related differential dynamics of EEG alpha– and beta-band synchronization in cortico-basal motor structures. Eur J Neurosci 25, 1604–1615 (2007).

44. Steina, A. et al. Mapping Subcortico-Cortical Coupling-A Comparison of Thalamic and Subthalamic Oscillations. Mov Disord 39, 684–693 (2024).

45. Litvak, V. et al. Resting oscillatory cortico-subthalamic connectivity in patients with Parkinson’s disease. Brain 134, 359–374 (2011).

46. Louie, K. H. et al. Cortico-Subthalamic Field Potentials Support Classification of the Natural Gait Cycle in Parkinson’s Disease and Reveal Individualized Spectral Signatures. eNeuro 9, ENEURO.0325-22.2022 (2022).

47. Buot, A. et al. Emotions Modulate Subthalamic Nucleus Activity: New Evidence in Obsessive-Compulsive Disorder and Parkinson’s Disease Patients. Biological Psychiatry: Cognitive Neuroscience and Neuroimaging (2020) doi:10.1016/j.bpsc.2020.08.002.

48. Zénon, A. et al. The human subthalamic nucleus encodes the subjective value of reward and the cost of effort during decision-making. Brain 139, 1830–1843 (2016).

49. Jensen, O. & Mazaheri, A. Shaping functional architecture by oscillatory alpha activity: gating by inhibition. Front Hum Neurosci 4, 186 (2010).

50. Lau, B. et al. The integrative role of the pedunculopontine nucleus in human gait. Brain 138, (2015).

51. He, S. et al. Gait-phase modulates alpha and beta oscillations in the pedunculopontine nucleus. Journal of Neuroscience (2021) doi:10.1523/JNEUROSCI.0770-21.2021.

52. Mori, S., Sakamoto, T., Ohta, Y., Takakusaki, K. & Matsuyama, K. Site-specific postural and locomotor changes evoked in awake, freely moving intact cats by stimulating the brainstem. Brain Res 505, 66–74 (1989).

53. Guillaumin, A., Serra, G. P., Georges, F. & Wallén-Mackenzie, Å. Experimental investigation into the role of the subthalamic nucleus (STN) in motor control using optogenetics in mice. Brain Res 1755, 147226 (2021).

54. Sanders, T. H. Stimulation of Cortico-Subthalamic Projections Amplifies Resting Motor Circuit Activity and Leads to Increased Locomotion in Dopamine-Depleted Mice. Front Integr Neurosci 11, 24 (2017).

55. Pan, M.-K. et al. Deranged NMDAergic cortico-subthalamic transmission underlies parkinsonian motor deficits. J Clin Invest 124, 4629–4641 (2014).

56. Amundsen Huffmaster, S. L., Lu, C., Tuite, P. J. & MacKinnon, C. D. The Transition from Standing to Walking Is Affected in People with Parkinson’s Disease and Freezing of Gait. J Parkinsons Dis 10, 233–243 (2020).

57. Oswal, A. et al. Neural signatures of hyperdirect pathway activity in Parkinson’s disease. Nature Communications (2021) doi:10.1038/s41467-021-25366-0.

58. Pardo-Valencia, J., Fernández-García, C., Alonso-Frech, F. & Foffani, G. Oscillatory vs. non-oscillatory subthalamic beta activity in Parkinson’s disease. J Physiol 602, 373–395 (2024).

59. Hall, L. M., Brauer, S. G., Horak, F. & Hodges, P. W. The effect of Parkinson’s disease and levodopa on adaptation of anticipatory postural adjustments. Neuroscience 250, 483–492 (2013).

60. Workman, C. D. & Thrasher, T. A. The influence of dopaminergic medication on balance automaticity in Parkinson’s disease. Gait Posture 70, 98–103 (2019).

61. Roytman, S. et al. Cholinergic system correlates of postural control changes in Parkinson’s disease freezers. Brain 146, 3243–3257 (2023).

62. Bohnen, N. I., Kanel, P., van Emde Boas, M., Roytman, S. & Kerber, K. A. Vestibular Sensory Conflict During Postural Control, Freezing of Gait, and Falls in Parkinson’s Disease. Mov Disord 37, 2257–2262 (2022).

63. Karachi, C. et al. Cholinergic mesencephalic neurons are involved in gait and postural disorders in Parkinson disease. J Clin Invest 120, 2745–2754 (2010).

64. Milekovic, T. et al. A spinal cord neuroprosthesis for locomotor deficits due to Parkinson’s disease. Nat Med 29, 2854–2865 (2023).

65. Cowie, D., Limousin, P., Peters, A. & Day, B. L. Insights into the neural control of locomotion from walking through doorways in Parkinson’s disease. Neuropsychologia 48, 2750–2757 (2010).

66. Delval, A. et al. Objective detection of subtle freezing of gait episodes in Parkinson’s disease. Mov Disord 25, 1684–1693 (2010).

67. Yin, Z. et al. Cortical phase-amplitude coupling is key to the occurrence and treatment of freezing of gait. Brain 145, 2407–2421 (2022).

68. Horn, A., Neumann, W.-J., Degen, K., Schneider, G.-H. & Kühn, A. A. Toward an electrophysiological ‘sweet spot’ for deep brain stimulation in the subthalamic nucleus. Hum Brain Mapp 38, 3377–3390 (2017).

69. Nambu, A. A new dynamic model of the cortico-basal ganglia loop. Prog Brain Res 143, 461–466 (2004).

70. Mink, J. W. The basal ganglia: focused selection and inhibition of competing motor programs. Prog Neurobiol 50, 381–425 (1996).

71. Schroll, H. & Hamker, F. H. Computational models of basal-ganglia pathway functions: focus on functional neuroanatomy. Front Syst Neurosci 7, 122 (2013).

72. Frank, M. J., Samanta, J., Moustafa, A. A. & Sherman, S. J. Hold your horses: impulsivity, deep brain stimulation, and medication in parkinsonism. Science 318, 1309–1312 (2007).

73. Aron, A. R. From reactive to proactive and selective control: developing a richer model for stopping inappropriate responses. Biol Psychiatry 69, e55–68 (2011).

74. Bouthour, W. et al. Biomarkers for closed-loop deep brain stimulation in Parkinson disease and beyond. Nat Rev Neurol 15, 343–352 (2019).

75. Basu, I. et al. Pathological tremor prediction using surface electromyogram and acceleration: potential use in ‘ON-OFF’ demand driven deep brain stimulator design. J Neural Eng 10, 036019 (2013).

76. Tan, H. et al. Decoding voluntary movements and postural tremor based on thalamic LFPs as a basis for closed-loop stimulation for essential tremor. Brain Stimul 12, 858–867 (2019).

77. Opri, E. et al. Chronic embedded cortico-thalamic closed-loop deep brain stimulation for the treatment of essential tremor. Sci Transl Med 12, eaay7680 (2020).

78. Ferleger, B. I. et al. Fully implanted adaptive deep brain stimulation in freely moving essential tremor patients. J Neural Eng 17, 056026 (2020).

79. He, S. et al. Closed-Loop Deep Brain Stimulation for Essential Tremor Based on Thalamic Local Field Potentials. Mov Disord 36, 863–873 (2021).

80. Melbourne, J. A. et al. Kinematic adaptive deep brain stimulation for gait impairment and freezing of gait in Parkinson’s disease. Brain Stimul 16, 1099–1101 (2023).

81. Velisar, A. et al. Dual threshold neural closed loop deep brain stimulation in Parkinson disease patients. Brain Stimul (2019) doi:10.1016/j.brs.2019.02.020.

82. He, S. et al. Beta-triggered adaptive deep brain stimulation during reaching movement in Parkinson’s disease. Brain 146, 5015–5030 (2023).

83. Oliveira, A. M. et al. Machine learning for adaptive deep brain stimulation in Parkinson’s disease: closing the loop. J Neurol 270, 5313–5326 (2023).

84. Busch, J. L. et al. Single threshold adaptive deep brain stimulation in Parkinson’s disease depends on parameter selection, movement state and controllability of subthalamic beta activity. Brain Stimul 17, 125–133 (2024).

85. O’Day, J. et al. The turning and barrier course reveals gait parameters for detecting freezing of gait and measuring the efficacy of deep brain stimulation. PLoS One 15, e0231984 (2020).

86. Strandquist, G. et al. Bringing the Clinic Home: An At-Home Multi-Modal Data Collection Ecosystem to Support Adaptive Deep Brain Stimulation. J Vis Exp (2023) doi:10.3791/65305.

87. Wilkins, K. B., Melbourne, J. A., Akella, P. & Bronte-Stewart, H. M. Unraveling the complexities of programming neural adaptive deep brain stimulation in Parkinson’s disease. Front Hum Neurosci 17, 1310393 (2023).

88. Schmidt, S. L. et al. At home adaptive dual target deep brain stimulation in Parkinson’s disease with proportional control. Brain 147, 911–922 (2024).

