## Supplemental material for "Subthalamic Signature of Freezing of Gait in Parkinson’s Disease"

**Supplementary Table 1.** Clinical data of the 38 patients included in the study

**Supplementary Table 2.** Demographic characteristics of the 23 control subjects

**Supplementary Fig. 1.** Parkinsonian motor disability and gait and balance disorders severity in Parkinson's disease patients

**Supplementary Fig. 2.** Gait initiation parameters and principal components analysis in Parkinson's disease patients

**Supplementary Fig. 3.** Local field potentials subthalamic neuronal activity in Parkinson's disease patients at gait initiation in FOG- and FOG+ patients, recorded in the DOPA$^{OFF}$ and DOPA$^{ON}$ conditions

**Supplementary Fig. 4.** Coordinates of the recordings dipoles of the posterior and central STN areas

**Supplementary Fig. 5.** Differences in STN neuronal activity links with gait initiation parameters between central and posterior STN subareas

**Supplementary Fig. 6.** Time-frequency maps of the correlation between STN-LFPs and gait initiation parameters in the posterior and central STN subareas, in FOG+ patients

**Supplemental Fig. 7** Time-frequency maps of the correlation between STN-LFPs and gait initiation parameters in the DOPA$^{ON}$ condition

### Supplemental Materials

| | Gender | Age | Disease duration (yrs) | LEED | UPDRS part III | | | Axial score | | | GABS score | | |
|---|---|---|---|---|---|---|---|---|---|---|---|---|---|
| | | | | | DOPA$^{OFF}$ | DOPA$^{ON}$ | % | DOPA$^{OFF}$ | DOPA$^{ON}$ | % | DOPA$^{OFF}$ | DOPA$^{ON}$ | % |
| **FOG+ patients** | | | | | | | | | | | | | |
| **GB_03** | F | 59 | 14 | 1000 | 53 | 27 | 49.1 | 11 | 5 | 54.5 | 18 | 10 | 44.4 |
| **GB_11** | M | 50 | 10 | 375 | 37 | 10 | 73.0 | 5 | 0 | 100 | 16 | 1 | 93.8 |
| **GB_14** | M | 51 | 26 | 1000 | 37 | 3 | 91.9 | 7 | 0 | 100 | 20 | 2 | 90 |
| **GB_18** | M | 68 | 11 | 1100 | 80 | 22 | 72.5 | 14 | 4 | 71.4 | _ | _ | _ |
| **GB_29** | M | 53 | 6 | 1200 | 28 | 4 | 75.7 | 5 | 1 | 80 | 16 | 4 | 75 |
| **GB_32** | F | 50 | 8 | 860 | 55 | 28 | 49.1 | 11 | 2 | 81.8 | _ | _ | _ |
| **GB_33** | M | 66 | 18 | 1175 | 34 | 6 | 82.3 | 6 | 1 | 83.3 | _ | _ | _ |
| **M_0101** | M | 54 | 8 | 1545 | 64 | 14 | 78.1 | 20 | 3 | 85 | 55 | 9 | 83.6 |
| **M_0103** | F | 51 | 12 | 2200 | 56 | 20 | 64.3 | 17 | 10 | 41.2 | 24 | 6 | 75 |
| **M_0203** | M | 62 | 9 | 860 | 44 | 2 | 95.5 | 12 | 0 | 100 | 23 | 0 | 100 |
| **M_0204** | M | 58 | 17 | 1725 | 35 | 11 | 68.6 | 15 | 7 | 53.3 | 43 | 12 | 72.1 |
| **M_0206** | M | 68 | 11 | 2200 | 23 | 6 | 73.9 | 6 | 1 | 83.3 | 17 | 1 | 94.1 |
| **Mean (SD)** | **9M/3F** | **57.5 (7.0)** | **12.5 (5.6)** | **1270 (551)** | **45.5 (16.5)** | **12.7 (9.3)*** | **72.8% (14.2)** | **10.8 (5.0)$** | **2.8 (3.1)*** | **77.8% (19.4)** | **25.7 (13.8)$** | **5.0 (4.4)*** | **80.9% (16.8)** |
| **FOG- patients** | | | | | | | | | | | | | |
| **GB_01** | M | 71 | 13 | 980 | 40 | 15 | 62.5 | 5 | 1 | 80 | 12 | 1 | 91.7 |
| **GB_02** | M | 63 | 10 | 2050 | 50 | 15 | 70 | 2 | 1 | 50 | 10 | 1 | 90 |
| **GB_04** | F | 57 | 10 | 1090 | 27 | 4 | 85.2 | 3 | 0 | 100 | 12 | 0 | 100 |
| **GB_05** | F | 67 | 12 | 1400 | 40 | 9 | 77.5 | 8 | 2 | 75 | 24 | 6 | 75 |
| **GB_06** | F | 66 | 13 | 850 | 53 | 16 | 69.8 | 15 | 4 | 73.3 | 12 | 0 | 100 |
| **GB_07** | F | 57 | 10 | 800 | 58 | 17 | 70.7 | 11 | 5 | 54.5 | 26 | 5 | 80.8 |
| **GB_09** | M | 60 | 11 | 1600 | 54 | 29 | 46.3 | 3 | 1 | 66.7 | 14 | 8 | 42.9 |
| **GB_10** | M | 57 | 8 | 1700 | 32 | 2 | 93.8 | 4 | 0 | 100 | 6 | 0 | 100 |
| **GB_12** | M | 51 | 15 | 1175 | 34 | 12 | 64.7 | 8 | 4 | 50 | 14 | 5 | 64.3 |

| | | | | | | | | | | | | | |
|---|---|---|---|---|---|---|---|---|---|---|---|---|---|
| **GB_13** | M | 40 | 8 | 1175 | 30 | 3 | 90 | 6 | 2 | 66.7 | _ | _ | _ |
| **GB_15** | F | 39 | 16 | 900 | 30 | 3 | 90 | 2 | 0 | 100 | _ | _ | _ |
| **GB_16** | M | 71 | 10 | 1000 | 26 | 5 | 80.8 | 4 | 3 | 25 | 8 | 7 | 12.5 |
| **GB_17** | M | 68 | 12 | 1000 | 37 | 7 | 81.1 | 7 | 2 | 71.4 | 12 | 1 | 91.7 |
| **GB_19** | M | 60 | 8 | 1000 | 33 | 9 | 72.7 | 4 | 3 | 25 | 6 | 2 | 66.7 |
| **GB_20** | F | 67 | 10 | 450 | 26 | 13 | 50 | 6 | 3 | 50 | 9 | 3 | 66.7 |
| **GB_22** | M | 29 | 13 | 1050 | 13 | 0 | 100 | 2 | 0 | 100 | 7 | 0 | 100 |
| **GB_23** | M | 61 | 8 | 950 | 28 | 4 | 85.7 | 5 | 1 | 80 | 9 | 1 | 88.9 |
| **GB_26** | F | 55 | 11 | 1400 | 30 | 2 | 93.3 | 4 | 0 | 100 | 9 | 0 | 100 |
| **GB_27** | F | 51 | 9 | 1500 | 11 | 0 | 100 | 8 | 1 | 87.5 | _ | _ | _ |
| **GB_28** | M | 50 | 12 | 1700 | 56 | 6 | 89.3 | 7 | 1 | 85.7 | _ | _ | _ |
| **GB_30** | M | 55 | 5 | 1000 | 57 | 2 | 96.5 | 4 | 0 | 100 | _ | _ | _ |
| **GB_31** | M | 52 | 8 | 750 | 30 | 6 | 80 | 3 | 0 | 100 | _ | _ | _ |
| **M_0102** | F | 63 | 11 | 800 | 34 | 7 | 79.4 | 8 | 2 | 75 | _ | _ | _ |
| **M_0201** | M | 60 | 13 | 1630 | 43 | 10 | 76.7 | 11 | 3 | 72.7 | 28 | 4 | 85.7 |
| **M_0202** | M | 63 | 16 | 875 | 51 | 10 | 80.4 | 5 | 2 | 60 | 11 | 0 | 100 |
| **M_0205** | M | 68 | 6 | 625 | 34 | 10 | 70.6 | 8 | 3 | 62.5 | 21 | 8 | 61.9 |
| **Mean (SD)** | **18M/8F** | **57.7 (10.2)** | **10.7 (2.8)** | **1133 (383)** | **36.8 (12.8)** | **8.3 (6.6)*** | **79.1% (13.8)** | **5.9 (3.2)** | **1.7 (1.5)*** | **73.5% (22.3)** | **13.2 (6.7)** | **2.7 (2.9)*** | **79.9% (23.3)** |
| **Total** | **27M/11F** | **57.6 (9.2)** | **11.3 (3.9)** | **1176 (439)** | **39.6 (14.4)** | **9.7 (7.7)*** | **77.1% (14.1)** | **7.4 (4.4)** | **2.1 (2.2)*** | **74.9% (21.3)** | **17.2 (12.0)** | **3.5 (3.5)*** | **80.2% (21.1)** |

**Supplementary Table 1. Clinical data of the 38 patients included in the study.** The table reports for each patient the gender, the age, the disease duration, the levodopa equivalent dose (LEED), the part III of the Unified Parkinson's Disease Rating Scale (UPDRS-III) and the Axial score (sum of the items "arising from chair", "gait", "freezing of gait", "postural stability" and "posture") without (DOPAOFF) and with (DOPAON) dopaminergic treatment, and the corresponding percentage of improvement. Asterisk indicates the significance of the paired t-test used to compare the DOPAOFF and the DOPAON conditions (FOG+: $t(11)=8.86$, $p<0.001$; FOG-: $t(25)=14.1$, $p<0.001$; Overall: $t(37)=16.80$, $p<0.001$). *$p<0.05$ compared to DOPAOFF condition (paired t-tests); \$$p<0.05$ between FOG+ and FOG- patients (Anova repeated measures and post-hoc Mann-Whitney U tests).

| # | Gender | Age (yrs) |
|---|---|---|
| **GB_FD_07** | M | 52 |
| **GB_RC_12** | F | 61 |
| **GB_FC_13** | F | 41 |
| **GB_BT_15** | M | 53 |
| **GB_AT_17** | F | 48 |
| **GB_BE_19** | M | 53 |
| **GB_CF_21** | F | 60 |
| **GB_BT_22** | F | 59 |
| **GB_DL_24** | F | 56 |
| **GB_PM_25** | F | 52 |
| **GB_DA_26** | M | 42 |
| **GB_DS_27** | F | 49 |
| **GB_MF_30** | M | 54 |
| **GB_FE_31** | M | 37 |
| **GB_DP_32** | M | 47 |
| **GB_SS_33** | F | 49 |
| **GB_BC_34** | F | 53 |
| **GB_GP_36** | M | 60 |
| **GB_BG_37** | M | 61 |
| **GB_CP_41** | M | 55 |
| **GB_LP_43** | M | 54 |
| **GB_BN_44** | M | 57 |
| **GB_FY_45** | M | 55 |
| **Mean (SD)** | **13M/10F** | **52.5 (6.4)** |

**Supplementary Table 2. Demographic characteristics of the 23 control subjects.** The table reports for each subject the gender and age.

**Supplementary results**

***Severity of gait and balance disorders, effects of dopaminergic medication and freezing predisposition***

FOG$^+$ patients had more severe gait and balance disorders compared to FOG$^-$ patients in the DOPA$^{OFF}$ condition (axial score FOG$^+$ - FOG$^-$ patients, estimate [SE]=4.86 [1.08], p<0.001; GABS score, estimate [SE]=12.62 [2.88], p<0.001, **Supplementary Fig. 1**) with no other significant demographic or clinical differences (**Supplementary Table S1**). Dopaminergic treatment significantly improved clinically measured parkinsonian motor signs and gait and balance disorders in all patients to a level that was not significantly different between FOG$^+$ and FOG$^-$ patients (axial score FOG$^+$ - FOG$^-$ patients, estimate [SE]=1.14 [1.08], p=1.00; GABS score, estimate [SE]=2.26 [2.88], p=0.44; **Supplementary Table 1, Supplementary Fig. 1**). We found a significant interaction between the effects of the dopaminergic medication on gait and balance symptom severity and FOG status (axial score x Group, $F_{(36)}$=14.39, p<0.001; GABS score x Group; $F_{(26)}$=10.41, p=0.003, ANOVA repeated measures, **Supplementary Table 1, Supplementary Fig. 1**).

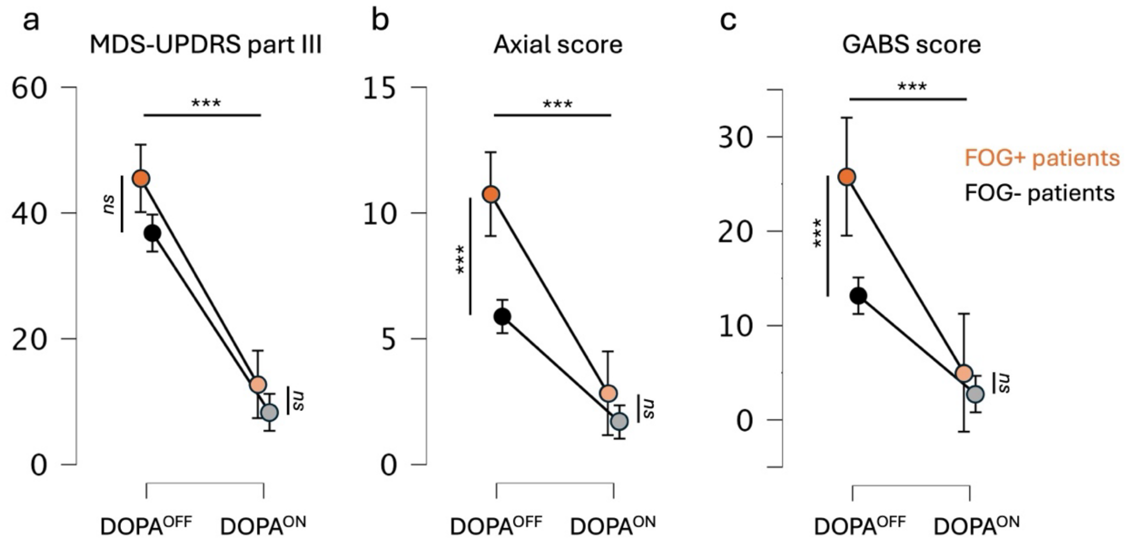


**Supplementary Fig. 1. Parkinsonian motor disability and gait and balance disorders severity in Parkinson's disease patients. a**, **b and c** Comparison of parkinsonian motor disability (MDS-UPDRS part III), axial and gait and balance scale (GABS) scores between patients with FOG during recordings (orange, FOG$^+$ patients, n=12 for MDRS UPDRS part III and axial scores, n=9 for GABS score) and those without FOG (black, FOG$^-$ patients, n=26 for MDRS UPDRS part III and axial scores, n=19 for GABS score, and between DOPA$^{OFF}$ and DOPA$^{ON}$ conditions. ***P<0.001, ANOVA repeated measures and Mann-Whitney post-hoc analysis, Bonferroni correctio). Each graph reports the mean and 95% confidence intervals.

***Link between pace, rhythm and balance in healthy controls***

In HC, we found significant relationship between pace (RPC1) and rhythm (RPC2) (r=-0.65, $t_{(2718)}$=-44.08, p<1.0e-04), pace and balance (RPC5) (r=0.17, $t_{(2718)}$=9.02, p<1.0e-04), and also between rhythm (RPC2) and balance (RPC5) (r=-0.19, $t_{(2718)}$=-10.07, p<1.0e-04).

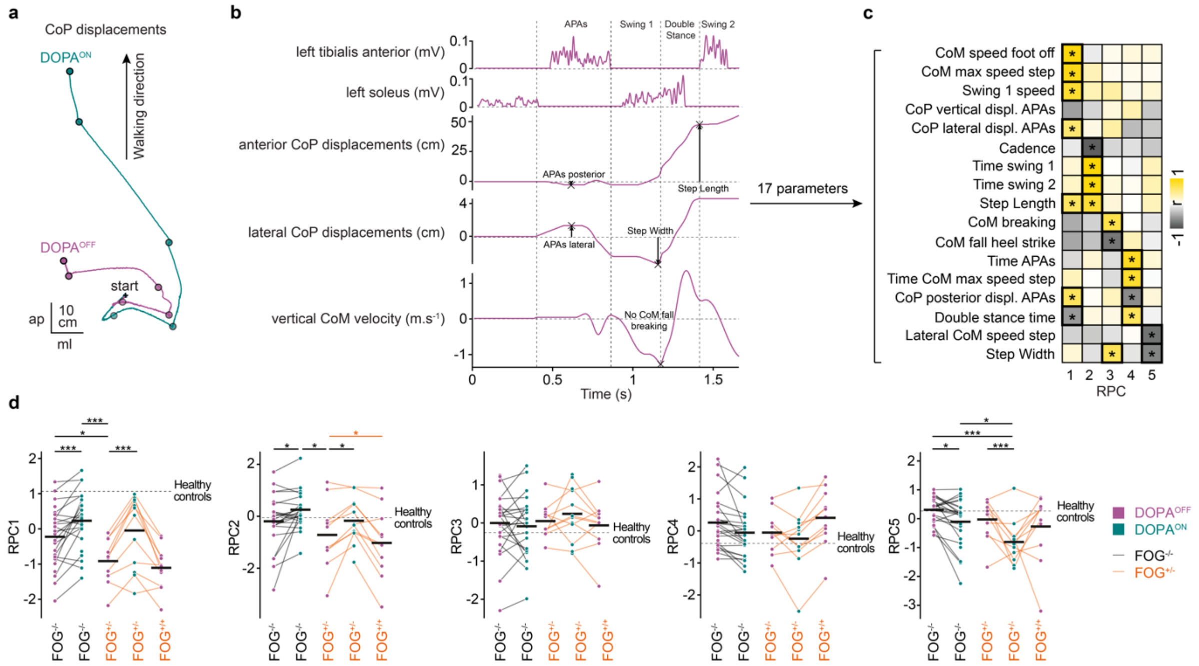


**Supplemental Fig 2. Gait initiation parameters and principal components analysis in Parkinson's disease patients. a** Gait initiation was recorded using a force plate, with calculation of the centre of foot pressure (CoP) anteroposterior and mediolateral displacements, and centre of mass (CoM) velocities in the 3 axis. **b.** Gait initiation includes two phases: 1) anticipatory postural adjustment phase (APAs), corresponding to the time between the first biomechanical event (APAs onset) and the 1$^{st}$ foot off and 2) the first step execution, corresponding to to the delay between the 1$^{st}$ foot off (start of the swing 1) and 1$^{st}$ foot contact (end of the swing 1 and start of the double stance period). **c**, 17 gait initiation parameters were included in the principal component analysis, with : CoP posterior and lateral displacements during the APAs, anterior CoM velocity at time of 1$^{st}$ foot-off (CoM speed foot off), maximal anterior velocity (CoM max speed step), anterior velocity at time of 1$^{st}$ step (swing 1 speed), step length, step width, minimal vertical velocities during the swing time (CoM fall heel strike) and at time of 1$^{st}$ foot contact and the differences between the two (CoM breaking), the cadence, swing time of the first and second steps (time swing 1 and time swing 2), duration of the APAs (time APA) and double stance (double stance time) phases, and mediolateral CoM velocity (Lateral CoM speed step) and time of the maximal CoM velocity (time CoM max speed step). **d**, RPCs scores obtained in patients without FOG episodes during recordings (FOG$^{-}$) in the DOPA$^{OFF}$ (purple circles) and DOPA$^{ON}$ (green circles) conditions, and in FOG+ patients, for FOG$^{+/-}$ trials in the DOPA$^{OFF}$ and DOPA$^{ON}$ conditions, and FOG$^{+/+}$ trials in the DOPA$^{OFF}$ condition. * $P < 0.05$, ** $P < 0.01$ and *** $P < 0.001$ for comparison between DOPA conditions and patient groups.

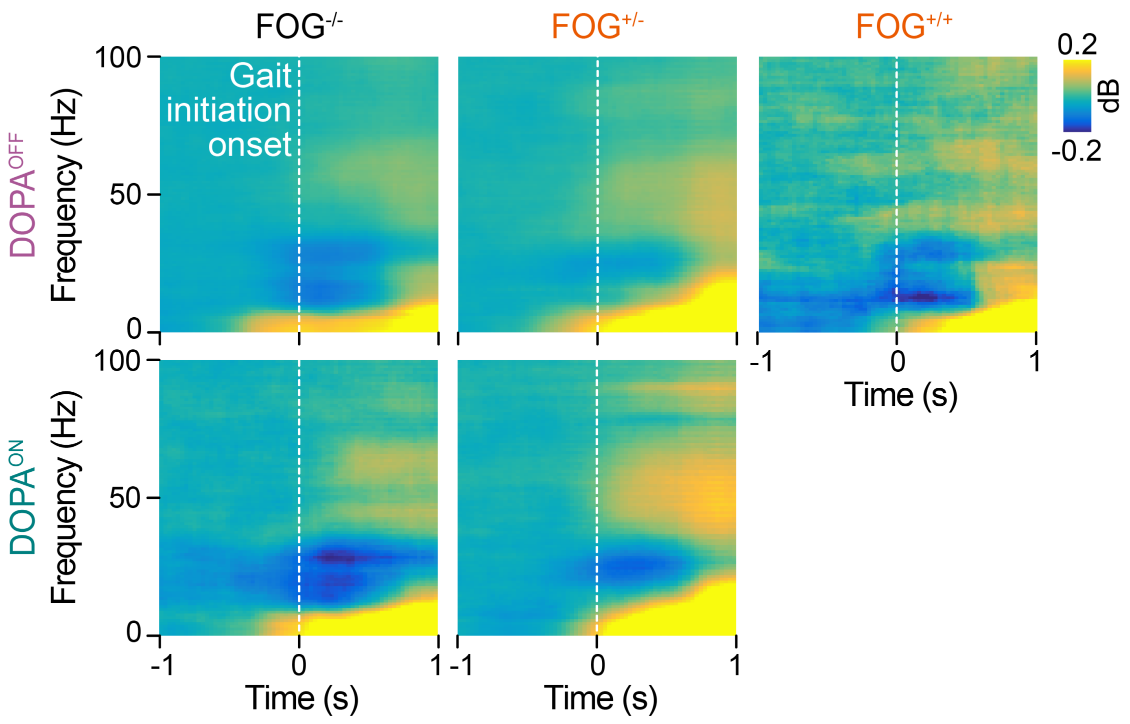


**Supplemental Fig. 3 STN LFPs at gait initiation in FOG$^-$ and FOG$^+$ patients in both DOPA$^{OFF}$ and DOPA$^{ON}$ conditions.** Time-frequency maps of the STN LFPs activity recorded in the DOPA$^{OFF}$ (upper panel) and DOPA$^{ON}$ (bottom panel) conditions, at gait initiation in FOG$^-$ (FOG$^{-/-}$ trials, left) and FOG$^+$ patients without (FOG$^{+/-}$ trials, middle) and with FOG (FOG+/+ trials, DOPA$^{OFF}$ condition) occurence.

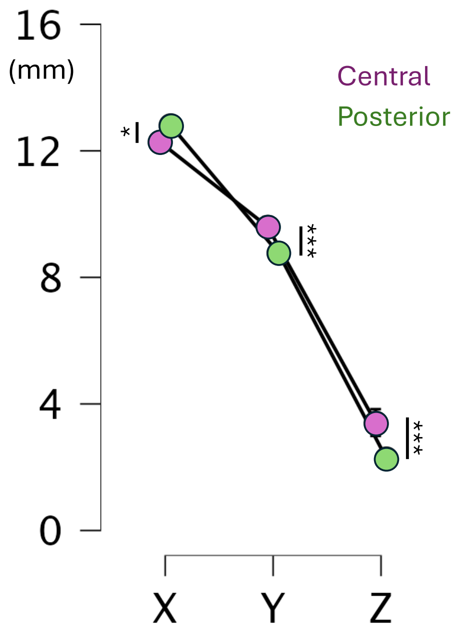


**Supplemental Fig. 4 Coordinates of the recordings dipoles of the posterior and central STN areas.**

Mediolateral (X), anteroposterior (Y) and depth (Z) coordinates of the recordings dipoles from the central (purple) and posterior (green) STN subareas. * $P < 0.05$, ** $P < 0.01$ and *** $P < 0.001$, using ANOVA repeated measures.

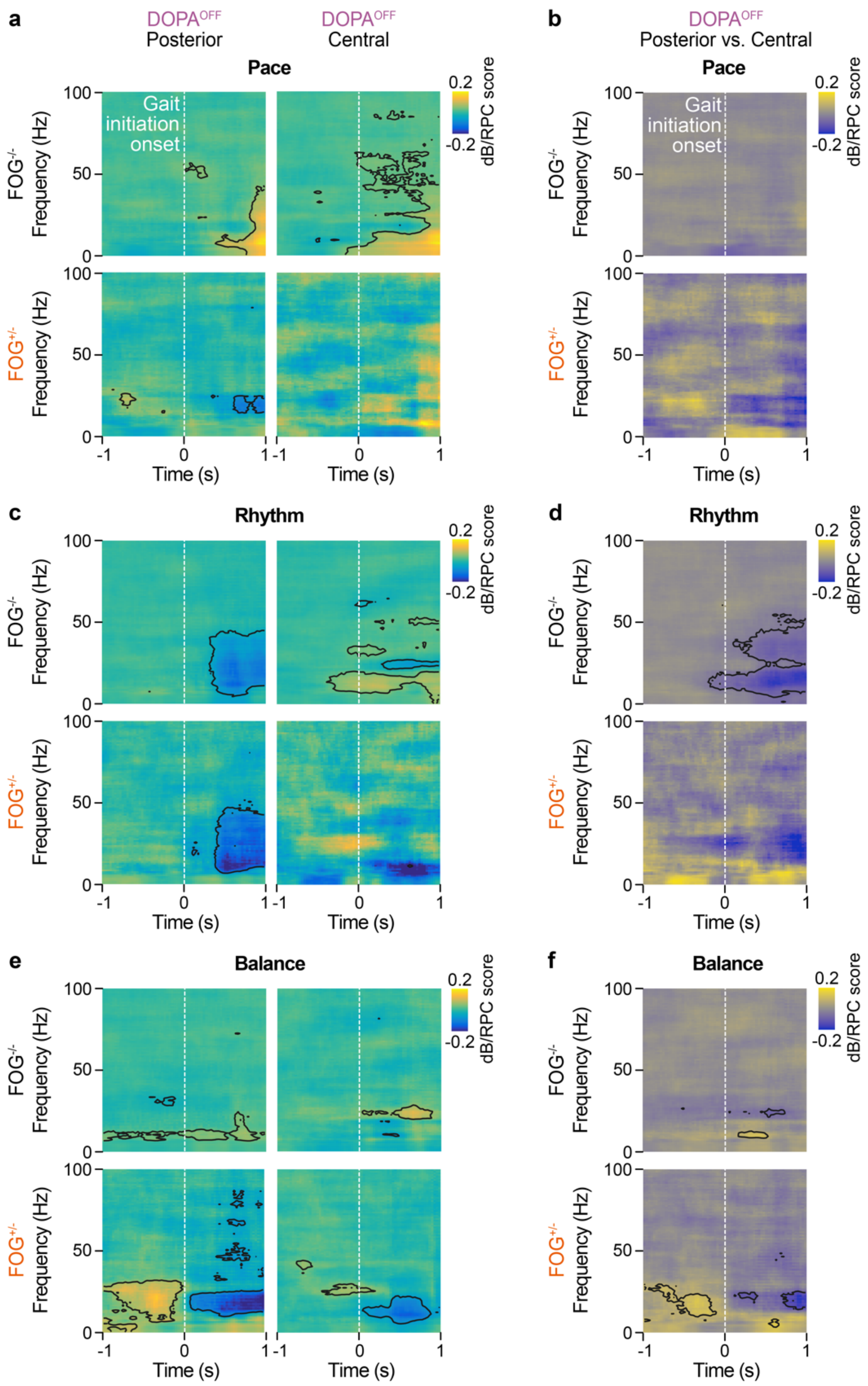


**Supplemental Fig. 5 STN neuronal activity modulation during gait initiation in the central and posterior STN subareas in FOG$^{-}$ and FOG$^{+}$ patients. a, b** Differences in TF maps associations between STN LFP recorded within the posterior and central STN and pace in FOG$^{-}$ (FOG$^{-/-}$ trials) and FOG$^{+}$ (FOG$^{+/-}$ trials) patients in the DOPA$^{OFF}$ condition. **c,d** same analysis for rhythm and **e,f** for balance scores. The black bars illustrate significant differences between the FOG$^{-/-}$ and FOG$^{+/-}$ trials.

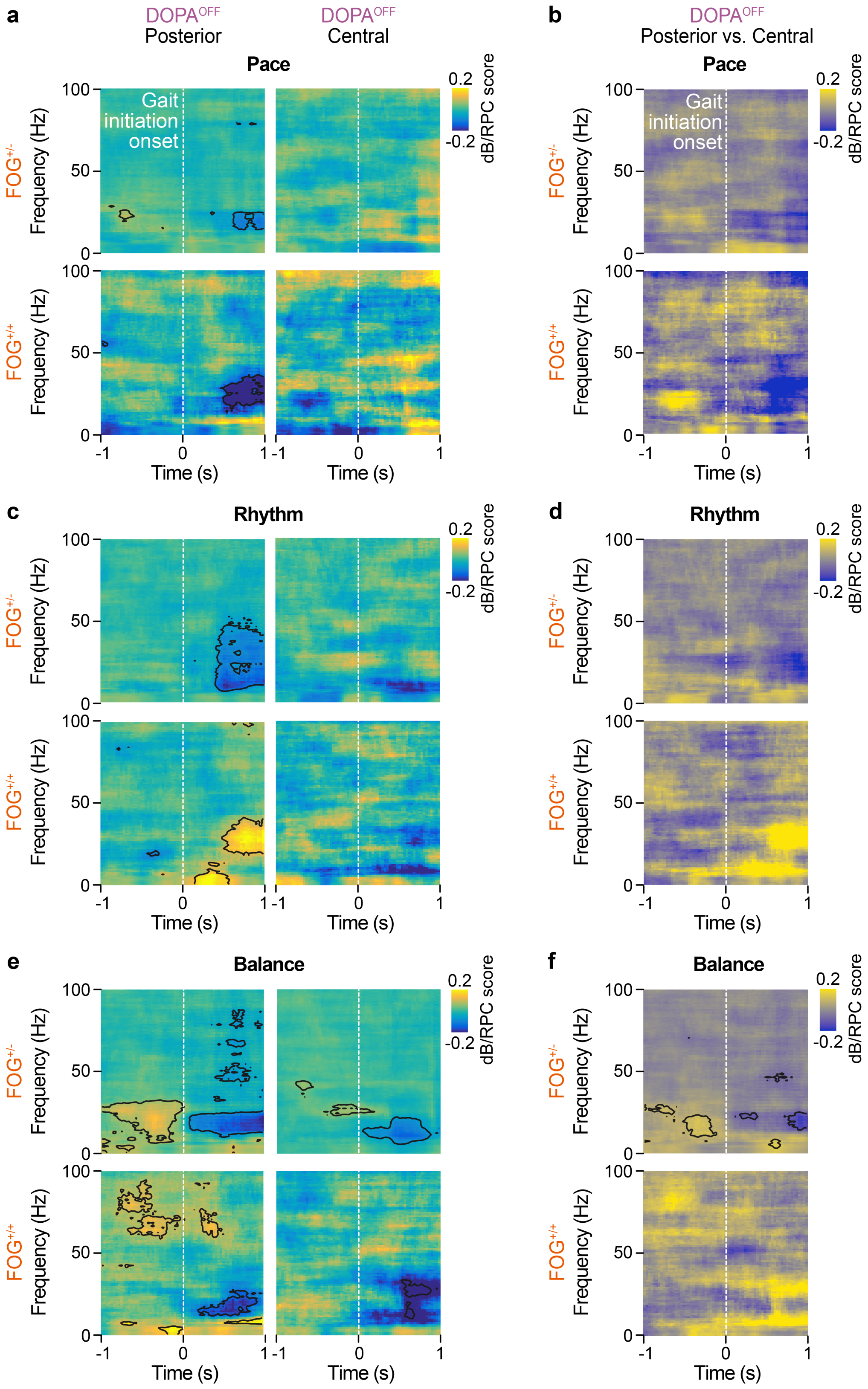


**Supplemental Fig. 6 STN neuronal activity modulation during gait initiation in the central and posterior STN subareas in FOG$^{+}$ patients for trials without and with FOG episodes. a, b** Differences in TF maps associations between STN LFP recorded within the posterior and central STN and pace for trials without (FOG$^{+/}$) and those with FOG episodes (FOG$^{+/+}$) in the DOPA$^{OFF}$ condition. **c,d** same analysis for rhythm and **e,f** for balance scores. The black bars illustrate significant correlation for both groups (**a, c and e**) and differences between the FOG$^{+/-}$ and FOG$^{+/+}$ trials (**b, d and f**).

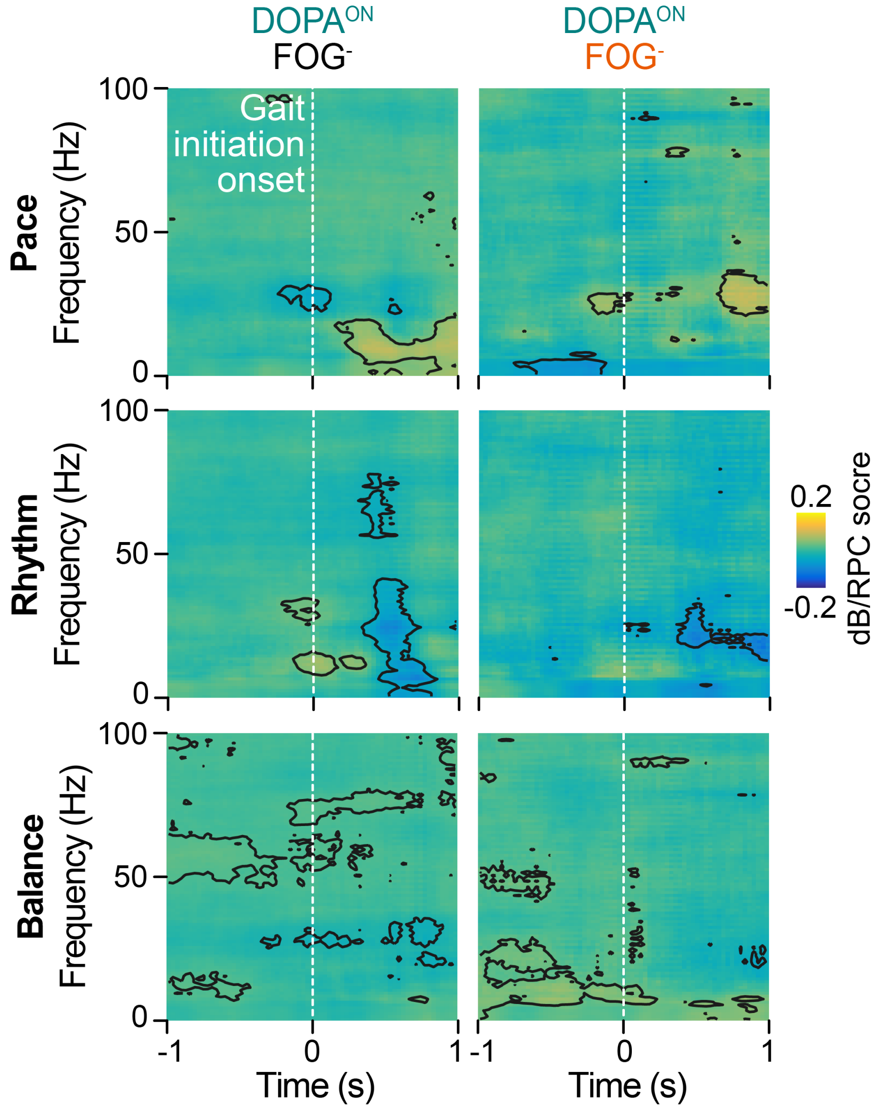


**Supplemental Fig. 7 Time-frequency maps of the correlation between STN-LFPs and gait initiation parameters in the DOPA$^{ON}$ condition**, for Pace, Rhythm and Balance scores, respectively, in FOG$^{-}$ and FOG$^{+}$ patients.
